# Mechanical Strain Activates Planar Cell Polarity Signaling to Coordinate Vascular Cell Dynamics

**DOI:** 10.1101/2024.06.25.600357

**Authors:** Lieke Golbach, Tanumoy Saha, Maria Odenthal-Schnittler, Jenny Lücking, Ana Velic, Emir Bora Akmeric, Dorothee Bornhorst, Oliver Popp, Philipp Mertins, Felix Gunawan, Holger Gerhardt, Boris Macek, Britta Trappmann, Hans J. Schnittler, Milos Galic, Maja Matis

**Author notes:** equal contribution, alphabetical order (Milos Galic, Maja Matis).

## Abstract

Mechanical stimuli, particularly laminar blood flow, play a crucial role in shaping the vascular system. Changes in the rate of blood flow manifest in altered shear stress, which activates signaling cascades that drive vascular remodeling. Consistently, dysregulation of the endothelial response to fluid shear forces and aberrant flow patterns both lead to pathological conditions, including impaired blood vessel development and atherosclerosis. Despite its importance, the mechanisms driving the coordinated cell behavior underlying vascular remodeling are not fully understood. Combining classical cell biological approaches with advanced image analysis, mathematical modeling, biomimetic strategies, and in vivo studies, we identify the planar cell polarity (PCP) protein Vangl1 as an enforcer of flow-dependent cell dynamics in the vascular system. We demonstrate that shear stress triggers the relocation of Vangl1 from an internal reservoir to the plasma membrane at the initiation of cell remodeling. Membrane enrichment of Vangl1 is mediated by a Coronin1C-dependent shift in the equilibrium between endo- and exocytosis and results in the spatial reorganization of another essential PCP protein, Frizzled6 (Fzd6). The resulting mutual exclusion of the core PCP proteins Fzd6 and Vangl1 augments differential junctional and cytoskeletal dynamics along the flow axis. Loss of Vangl1 limits the ability of endothelial cells to respond to shear forces in a coordinated fashion, resulting in irregular cell alignment along the flow direction and erroneous vessel sprouting. Together, these studies introduce core PCP signaling as a determinant of collective cell dynamics and organization of the vascular system.

## INTRODUCTION

The detection and accurate interpretation of mechanical stimuli is central to the development and function of organs at the cellular and tissue scale. One prominent example is the endothelium, the innermost layer of the vascular system, which translates frictional forces from blood flow (i.e., fluid shear stress) into distinct biochemical signals that control cell behavior (Hahn and Schwartz, 2009; Tanaka et al., 2021; Yamamoto and Ando, 2011). These pathways shape the vascular system not only during development but also provide continuous flow-dependent adjustments throughout life.

Under physiological conditions, laminar shear stress triggers the elongation of endothelial cells and polarized distribution of cytoskeleton and organelles along the axis of the blood flow (Dieterich et al., 2000; Franke et al., 1984; McCue et al., 2006; Noria et al., 1999; Noria et al., 2004; Tzima et al., 2003). This sensitivity of endothelial cells to shear stress is critically involved in several developmental and physiological vascular processes, such as angiogenesis and vascular morphogenesis, as well as vascular remodeling (Bondareva et al., 2019; Chen and Tzima, 2009; Franco et al., 2015; Poduri et al., 2017). Consistently, impairment of either the forces or the mechanisms by which these forces are sensed yield vascular disease (Chiu and Chien, 2011; Hahn and Schwartz, 2009). A coherent molecular mechanism of flow sensing and global coordination of individual cells within the vasculature, however, is currently still lacking.

One of the principal signaling pathways that instruct individual cells to generate tissue-scale rearrangements within the plane of a cell sheet is the planar cell polarity (PCP) signaling pathway (Butler and Wallingford, 2017; Matis and Axelrod, 2013). PCP is established by an evolutionarily conserved set of proteins that assemble within cells in mutually exclusive complexes, leading to the coordinated polarization of downstream targets within each cell of the planarly polarized tissue (Galic and Matis, 2015). Importantly, PCP signaling not only directs the orientation of subcellular structures in non-motile epithelial cells but also coordinates dynamic cell behavior within the plane (Matis and Axelrod, 2013). As such, PCP plays a central role in the orientation of cell divisions, tissue growth, cell rearrangements, cell migration and differentiation (Aigouy et al., 2010; Baena-Lopez et al., 2005; Blair and McNeill, 2018; Goodrich and Strutt, 2011; Mao et al., 2016; Mao et al., 2011; Zakaria et al., 2014). Consistently, mutations in PCP genes contribute to developmental anomalies and diseases (Badouel et al., 2015; Copp and Greene, 2010; Ebnet et al., 2018; Ishiuchi et al., 2009; Simons et al., 2005; Simons and Mlodzik, 2008; Zakaria et al., 2014).

Unlike the role of PCP signaling in epithelial tissue, which is well established, comparatively little is known about its function in endothelial cells. In support of the role of PCP signaling during vascularization, loss-of-function studies in mice showed that Frizzled4 (Fzd4) and Frizzled6 (Fzd6) affect the patterning of arterial vessel morphogenesis (Descamps et al., 2012; Markovic et al., 2017), while Celsr1 together with Vangl2 regulates directed cell rearrangements during lymphatic valve formation (Tatin et al., 2013). However, while these results establish the presence and function of some PCP proteins in endothelial cells, the dynamics of antagonistic PCP complexes and the functional importance of mechanical stress for PCP signaling in the vascular system remain both unclear. Here, we examine how core PCP proteins respond to shear forces, and whether they contribute to global patterning of the endothelium. We find that fluid shear stress activates polarization of the core PCP module to regulate tissue-scale remodeling of the vasculature.

## RESULTS

### Vascular remodeling depends on core PCP proteins

At the molecular level, PCP-dependent signaling relies on six proteins. These include three multipass transmembrane proteins, Frizzled (Fzd) (Vinson et al., 1989) Van Gogh (Vangl) (Taylor et al., 1998; Wolff and Rubin, 1998), Flamingo (Celsr) (Usui et al., 1999), and the cytoplasmic proteins Dishevelled (Dvl) (Klingensmith et al., 1994; Theisen et al., 1994), Prickle (Pk) (Gubb et al., 1999) and Diego (Dg) (Feiguin et al., 2001). PCP is established through the mutually exclusive polarization of the core PCP complexes Fzd-Dvl and Vangl-Pk (Butler and Wallingford, 2017). Probing the expression of core PCP genes in human umbilical vein endothelial cells (HUVECs), we found high transcript levels of Fzd6, Dvl1-3, Vangl1 and Pk1 (**Fig. S1**), establishing the presence of both core PCP complexes in HUVECs.

Having identified core PCP proteins (**Fig. S3**), we next aimed to determine their role. Endothelial cells experience a range of shear stress profiles that differ in magnitude, direction, and temporal characteristics depending on their location within the vasculature. In addition to spatial differences, shear stress at a particular site can also change over time, causing local endothelial remodeling (Baeyens et al., 2014; Levesque and Nerem, 1985; Wang et al., 2012). For instance, cultured HUVECs respond to shear forces of 10-20 dyn/cm^2^, which corresponds to the physiological flow within the umbilical vein (Boito et al., 2002; Kiserud and Rasmussen, 1998) by aligning along the flow axis (Baeyens et al., 2015). To explore the mechanisms that guide this long-range planar polarization of the endothelium, we first established the collective behavior of cells exposed to laminar shear forces and compared them to cells cultured without flow. Throughout all experiments, HUVECs were plated and grown to a density of 600 cells/mm^2^ and then measured for the flow-dependent response using custom-made software (**Fig. S2a,b**). As previously published (Baeyens et al., 2015), we found a rapid onset of cell elongation and alignment along the flow axis within the first 12 hours upon exposure to a constant flow of 18 dyn/cm^2^ (**Fig. 1a**). In comparison, cells plated without flow showed no changes in cell shape (**Fig. S2c-g**), consistent with the established role of flow for the collective behavior of endothelial cells. The inability of cells to align was further associated with a reduction in mean speed (**Fig. S2f-g**).

**Figure 1:**
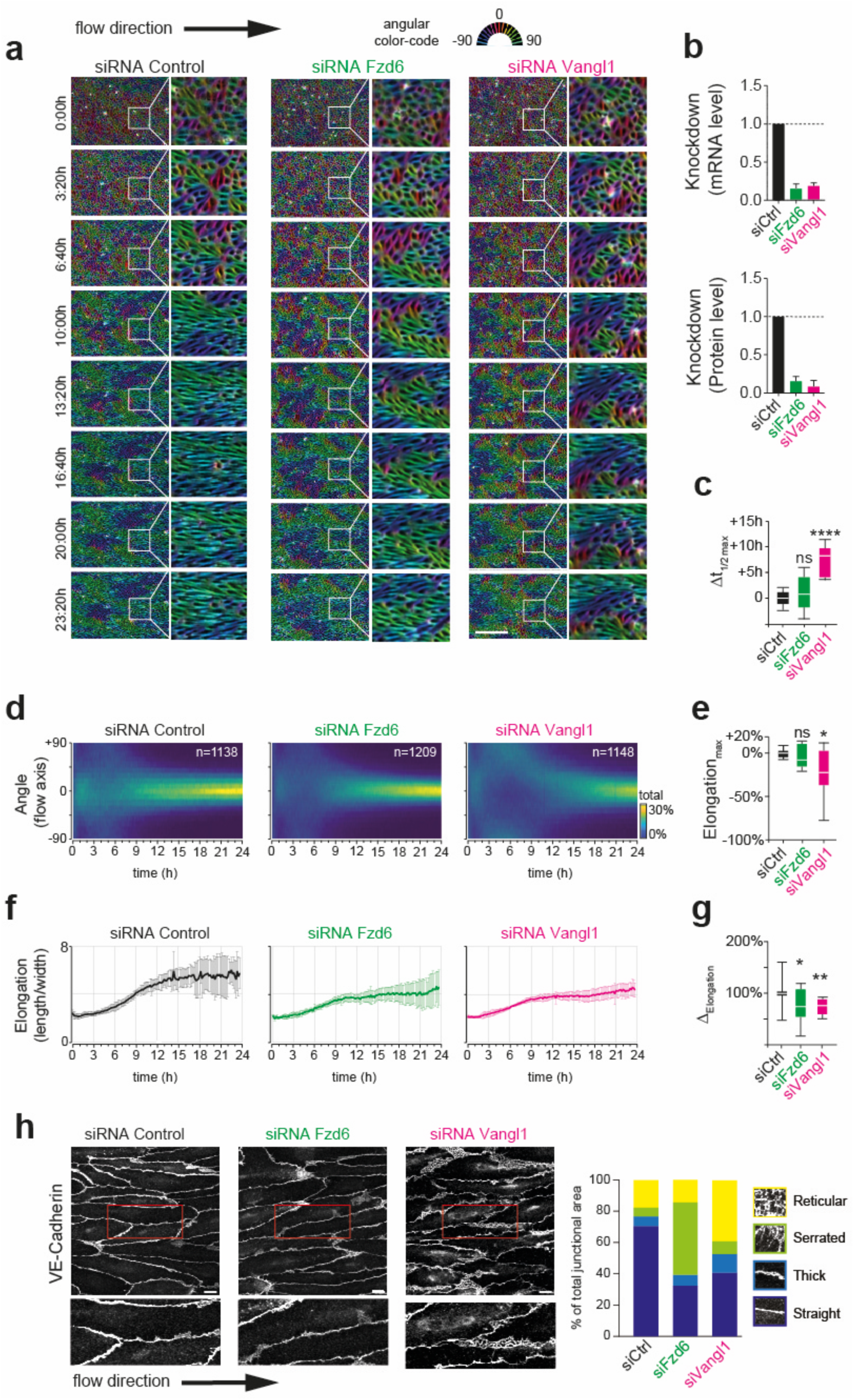
Planar cell polarity proteins regulate shape changes and collective cell dynamics along the flow axis. **(a)** Compared to control cells (left), the knockdown of core PCP proteins Fzd6 (middle) and Vangl1 (right) weakens cell elongation along the flow axis. **(b)** Validation of knockdown levels at the transcriptional (top) and protein (bottom) levels normalized to control (siCtrl). (siRNA Control: N = 3 biological repeats; siRNA Fzd6: N = 3 biological repeats; siRNA Vangl1: N = 3 biological repeats). For all plots, the control is black, Fzd6 is green, and Vangl1 is magenta. **(c)** Quantification of shape changes shows a significant delay of cell elongation upon knockdown of Vangl1 (blue). (Ordinary one-way ANOVA: n.s. p = 0.6234, **** p < 0.0001, siRNA Control: N = 3 biological repeats, n = 1138 cells; siRNA Fzd6: N = 3 biological repeats, n = 1209 cells; siRNA Vangl1: N = 3 biological repeats, n = 1148 cells). **(d)** Angular distribution of major cell axis over time. **(e)** Quantification of angular distribution shows a significant reduction in the alignment of the cells along the flow axis upon knockdown of Vangl1 (blue). (Ordinary one-way ANOVA: n.s. p = 0.9412, * p = 0.0294, siRNA Control: N = 3 biological repeats, n = 1138 cells; siRNA Fzd6: N = 3 biological repeats, n = 1209 cells; siRNA Vangl1: N = 3 biological repeats, n = 1148 cells). **(f)** Quantification of cell elongation (length/width) over time. (siRNA Control: N = 3 biological repeats, n = 1138 cells; siRNA Fzd6: N = 3 biological repeats, n = 1209 cells; siRNA Vangl1: N = 3 biological repeats, n = 1148 cells). **(g)** Quantification of cell length shows a significant reduction upon knockdown of Vangl1 (blue). (Ordinary one-way ANOVA: * p = 0.0461 and ** p = 0.0096, siRNA Control: N = 3 biological repeats, n = 1138 cells; siRNA Fzd6: N = 3 biological repeats, n = 1209 cells; siRNA Vangl1: N = 3 biological repeats, n = 1148 cells). **(h)** Analysis of junctional morphology (marked by VE-cadherin) shows changes in cell shape upon loss of Fzd6 and Vangl1 (siRNA Control: N = 5 biological repeats, n = 154 cells; siRNA Fzd6: N = 3 biological repeats, n = 93 cells; siRNA Vangl1: N = 3 biological repeats, n = 60 cells). Scale bars, (**a**) 100 μm, (**h**) 10 μm.

To study the functional relevance of both core PCP complexes, we separately targeted Vangl1 and Fzd6 via siRNA. qPCR and Western blotting showed high knockdown efficiency at the RNA and protein levels, respectively, for both constructs (**Fig. 1b** and **Fig. S3**). Next, we probed for functional changes upon loss of Vangl1 and Fzd6, respectively. We find that the depletion of Vangl1 or Fzd6 diminished the response of HUVECs to shear forces. Specifically, we find a significant delay in cell elongation (**Fig. 1a-c**), as well as a reduction in the ability of cells to align along the flow axis (**Fig. 1d-e**). For all parameters, we observed a stronger effect upon loss of Vangl1 compared to Fzd6.

We next sought to identify the mechanisms by which Vangl1 transduces shear forces into the endothelial cells. In epithelial cells, core PCP proteins direct cell behavior through the polarization of cytoskeletal elements and cell adhesion (Butler and Wallingford, 2017; Davey and Moens, 2017). Therefore, we analyzed its effects on the adherens junction complex, a key mediator of junctional remodeling during endothelial cell rearrangements (Cao et al., 2017; Huveneers et al., 2012). Crucial for junctional remodeling is the continued ability of cell-cell junctions to assemble, rearrange and disassemble, which requires a transition between actively remodeling and stable, mature junctions (Huveneers et al., 2012; Ngok et al., 2012). To probe for possible junctional changes upon Vangl1and Fzd6 knockdown, we binned the cell boundary into four previously defined morphological categories: serrated junctions (Bentley et al., 2014; Huveneers et al., 2012), reticular junctions (Fernandez-Martin et al., 2012; Huveneers et al., 2012) and JAILs (i.e., Junction-Associated Intermediate Lamellipodia) (Cao et al., 2017), thick junctions and straight junctions (Huveneers et al., 2012) (**Fig. 1h**). Control cells displayed straight junctions after 24 hours of flow exposure, while loss of Vangl1 led to a significant decrease of these junctions and a concomitant increase of reticular junctions. In contrast, Fzd6 knockdown increased the frequency of serrated junctions, establishing distinct effects on junctional stability for the two core PCP proteins (**Fig. 1h**). Together, the observed changes in cell shape and junctional morphology suggest that Vangl1 and Fzd6 contribute to polarized junctional dynamics.

### Core PCP proteins Fzd6 and Vangl1 arrive sequentially at the cell surface

In cells expressing PCP proteins, reports indicate their localization to the plasma membrane; however, the pattern varies, and consistent segregation of different PCP complexes is not uniformly documented (Davey and Moens, 2017). To determine the distribution of the key PCP components, Vangl1 and Fzd6, we performed immunocytochemistry. Using cells that were exposed for different intervals of time to shear forces and subsequently stained with antibodies directed against endogenously expressed Fzd6 and Vangl1, we find Fzd6 to be continuously present at the cell membrane (**Fig. 2a,b**). In contrast, Vangl1 signal intensity at the plasma membrane increased only after exposure to laminar flow (**Fig. 2a,b**). The obtained result contrasts with the behavior of core PCP proteins in other systems, especially in epithelial cells, where they localize to the membrane immediately after their synthesis.

**Figure 2:**
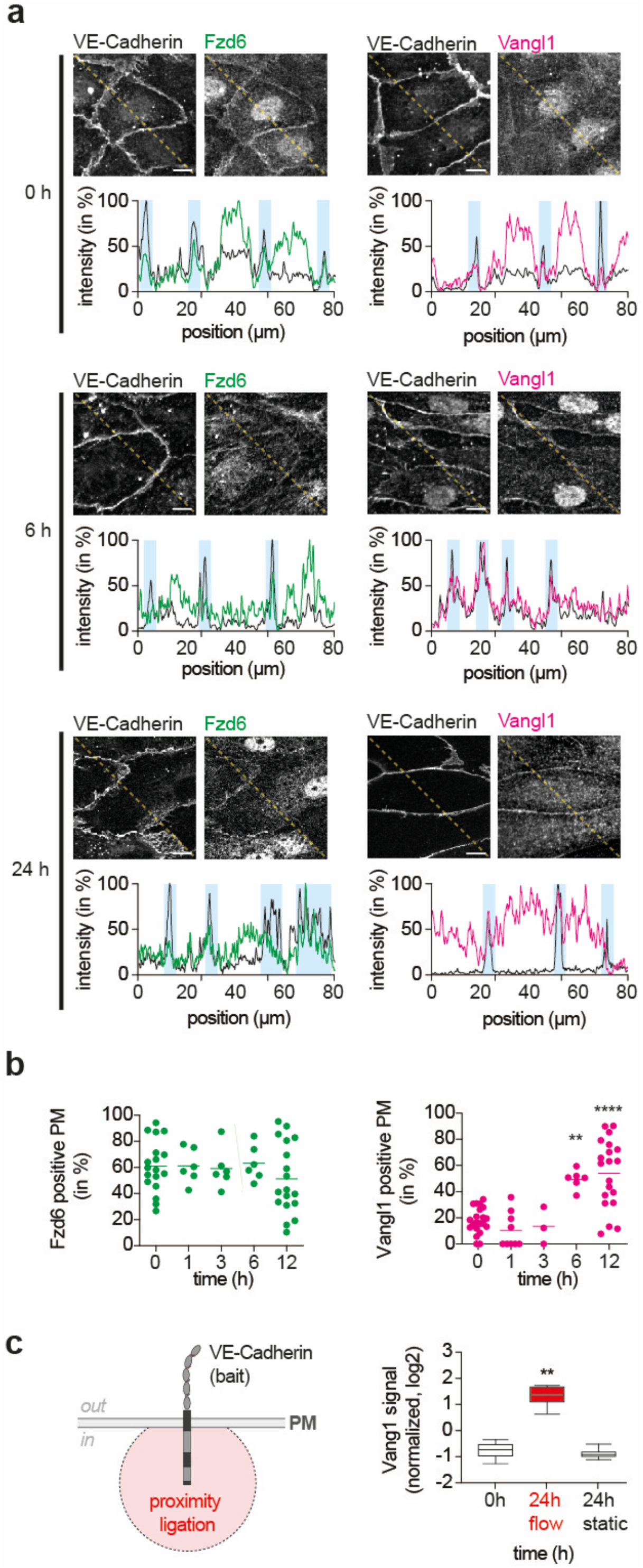
Vangl1 shows delayed membrane recruitment upon flow induction. **(a)** Top: Immunocytochemistry shows Fzd6 is continuously present at the cell junctions (marked by VE-cadherin), while Vangl1 only enriches at the cell junctions upon exposure to laminar flow. Below is the intensity profile of the yellow dotted line in the image above the blot (VE-cadherin black, Fzd6 green, and Vangl1 magenta). **(b)** Quantification of Fzd6 (green) and Vangl1 (magenta) enrichment at the plasma membrane. (0h, N = 18 repeats; 1h, N = 6 repeats; 3h, N = 6 repeats; 6h, N = 6 repeats; 12h, N = 18 repeats). **(c)** Proximity ligation, using VE-Cadherin as bait in the presence and absence of flow. For Vangl1, enrichment at the cell surface only upon flow induction is detected (t-test: ** p=?, 0h, N = 4 repeats; 24h flow, N = 6 repeats; 24h static, N = 4 repeats). Scale bar, (a) 10 μm.

To confirm these findings, we performed proximity ligation before and after the flow induction using VE-Cadherin as bait. Specifically, we fused the biotin ligase TurboID, which promiscuously labels proteins with a labeling radius of ∼10 nm (Branon et al., 2018), to the intracellular domain of VE-Cadherin and expressed it in HUVECs in the presence and absence of shear forces for 24 hours (**Fig. 2c, left** and **Fig. S4a**). Consistent with our findings, subsequent MS analysis of the proteins located proximal to VE-Cadherin identified Vangl1 only upon flow induction (**Fig. 2c, right**). The same trend was observed for Prickle1 (**Fig. S4b**), which forms a complex with the Vangl1. However, pulldown experiments using Vangl1 as bait failed to elute VE-Cadherin (**Fig. S4c**), arguing against direct interactions between these two proteins at the cell surface.

### Coronin1C mediates delayed surface localization of Vangl1

The inducible increase of surface Vangl1 levels establishes that shear forces control PCP protein dynamics at the plasma membrane. To determine whether this dynamic occurs at the transcriptional level, we compared the total protein amount before and after exposure to laminar shear stress. Intriguingly, quantification showed that Vangl1 protein levels stayed the same. Thus, the transcriptional regulation of protein cannot explain the observed increase of surface-localized Vangl1 (**Fig. 3a** and **Fig. S5**). Rather, in the absence of fluid shear forces, we find Vangl1 to be localized in intracellular vesicles (**Fig. 3b**), suggesting regulation of protein localization instead of protein levels. Since Vangl is subjected to membrane turnover in other cellular systems (Cho et al., 2015; Strutt and Strutt, 2008; Strutt et al., 2011), we inhibited endocytosis and probed for changes in Vang1 localization. The addition of the Dynasore, which blocks Dynamin-dependent vesicle scission (Macia et al., 2006), significantly increased Vangl1 levels at the plasma membrane (**Fig. 3c**).

**Figure 3:**
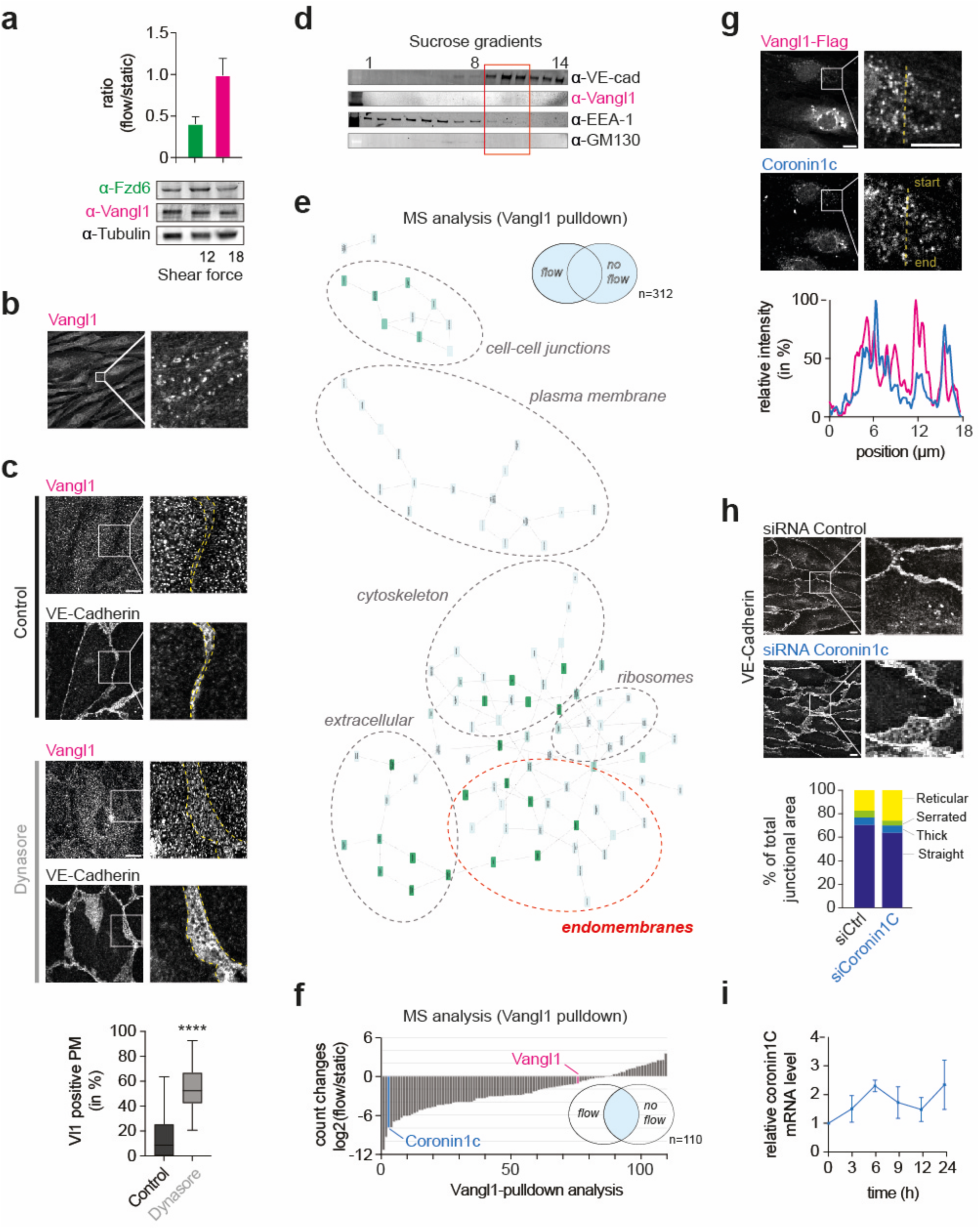
Coronin1C regulates the transport of Vangl1 to the cell surface. **(a)** Western blot analysis of Fzd6 and Vangl1 levels upon exposure to shear forces. Note stable expression levels for Vangl1, while Fzd6 levels are lowered upon flow induction. (N = 3 repeats). **(b)** Immunocytochemistry of cells shows Vangl1 localization in intracellular vesicles before induction of flow. **(c)** Global inhibition of endocytosis through Dynasore augments Vangl1 at the surface. (Mann-Whitney test: **** p < 0.0001, Control: n = 58 cells; Dynasore: n = 49 cells). **(d)** Cell fractionation shows Vangl1 localization to distinct endocytic vesicles. **(e)** Network representation of cellular components identified in Vangl1 pulldown of whole cell lysates using GOnet (https://tools.dice-database.org). (N = 3 static and 2 with flow). **(f)** MS analysis of cells using Vangl1 as bait identifies Coronin1C as a potential target. Proteins were ranked based on iBAQ counts. (N = 3 static and 2 with flow). **(g)** Vangl1 and Coronin1C co-localize in endocytotic vesicles. Below, the intensity profile of the yellow dotted line from the image are shown (Coronin1c blue and Vangl1 magenta). **(h)** Analysis of junctional morphology upon loss of Coronin1C. **(i)** Corinin1C increased expression levels coincide with the increase in surface Vangl1 localization. (N = 3 repeats). Scale bars, (b,c,g,h) 10 μm.

Having established endocytic trafficking of Vangl1, we next aimed to understand how this is accomplished. To probe for factors that reside with Vangl1 at endosomes and potentially regulate its transport to the cell surface, we conducted subcellular membrane vesicle fractionation using density gradient ultracentrifugation. We found that neither EEA-1, a marker of early endosomal vesicles, nor GM130, a Golgi marker, colocalized with Vangl1 (**Fig. 3d**), suggesting its localization to distinct endocytic vesicles. To quantify this observation more comprehensively, we conducted mass spectrometry (MS) analysis on the fraction containing Vangl1. Consistent with the notion that Vang1 resides in intracellular vesicles, gene ontology (GO) analysis showed enrichment of endo-membranes (**Fig. S6a-c**). In a complementary approach, we then performed an MS analysis of a pulldown assay with Vangl1 as bait. Since shear forces mediate the relocalization of Vangl1 from intracellular vesicles to the plasma membrane, pulldowns for both conditions were probed. GO analysis showed enrichment for proteins annotated to endomembranes (**Fig. 3e** and **Fig. S6**). Considering that interaction partners of Vangl1 may differ based on its subcellular localization, we ranked all proteins present in both conditions. Among the three proteins with the strongest change, we found Actin (ACTG1), Myosin10 (MYH10), and Coronin1C (CORO1C) (**Fig. 3f** and **List S1**).

Coronins play essential roles in regulating the actin cytoskeleton by interacting with actin filaments and other actin-binding proteins (Chan et al., 2011). Depending on the cellular context and availability of interacting partners, Coronins act as positive or negative regulators. However, a growing body of work supports additional functions in endocytosis (Kimura et al., 2008), endosomal fission (Hoyer et al., 2018) and exocytosis (Yi et al., 2002). Considering the gating properties of Coronin1C for endocytosed proteins back to the plasma membrane, we probed for a functional connection to Vangl1. Intriguingly, Vangl1 and Coronin1C partially co-localized in vesicles (**Fig. 3g**). Next, we knocked down Coronin1C and quantified changes in junctional morphology. Similar to cells lacking Vangl1 (**Fig. 1h**), we find increased formation of reticular junction upon loss of Coronin1C (**Fig. 3h**). Notably, we further observed Coronin1C levels to increase concomitantly with Vangl1 surface localization (**Fig. 3i**), reinforcing the notion that Coronin1C may regulate the subcellular Vangl1 localization in endothelial cells.

### Fzd6 and Vangl1 were enriched at opposite sides of the cell

Having established the effects of core PCP proteins on junctional morphology, we next probed its subcellular localization. Cells were subjected to shear forces for 3 or 12 hours and then fixed. Next, cells were co-stained with antibodies directed against VE-Cadherin and either Fzd6 or Vangl1. After three hours of flow, Fzd6 displayed a uniform distribution but became enriched at serrated junctions after 12 hours of flow (**Fig. 4a, top**). In contrast, Vangl1 was enriched at stable junctions from the beginning (**Fig. 4a, bottom**), suggesting that the junctional localization of Vangl1 limits the distribution of Fzd6 at the plasma membrane. To test this possibility, we next co-stained cells with antibodies directed against both core PCP proteins. Consistent with our previous findings, we find Fzd6 and Vangl1 to form mutually exclusive regions within single cells along the longest axis (**Fig. 4b** and **Fig. S8a**). Strikingly, the polarized accumulation of PCP proteins provides a possible explanation for the knockdown-dependent changes in endothelial remodeling described above. The loss of junctional polarity of Fzd6 in Vangl1 knockdown cells, which yields a spread of active reticular junctions (**Fig. 1h**), suggests that a flow-induced polarization of Fzd6 through Vangl1 is required for limiting junctional dynamics to the cell front.

**Figure 4:**
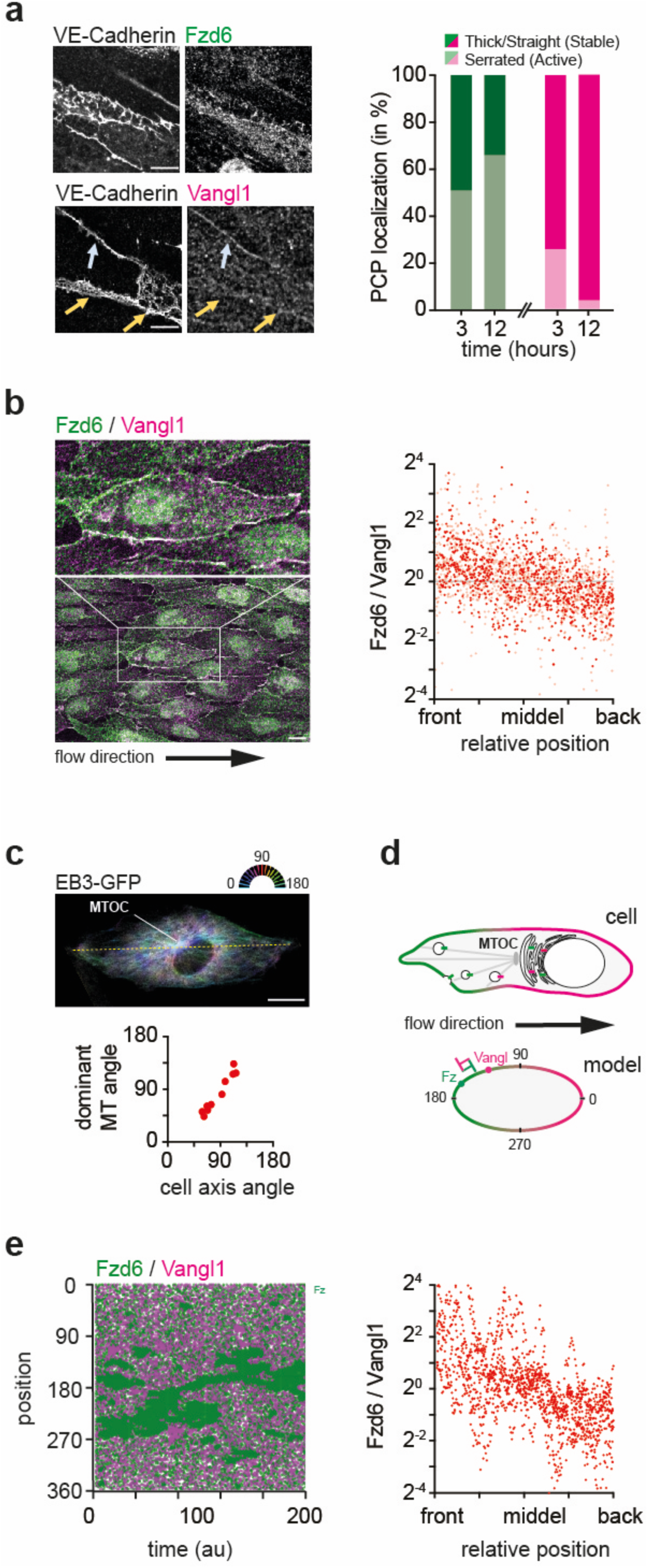
Core PCP proteins Fzd6 and Vangl1 polarize along the flow axis. **(a)** Vangl1 and Fzd6 enrich at cell junctions (marked with VE-cadherin) with distinct morphologies. Note increased accumulation of Fzd6 (green) at serrated junctions and of Vangl1 (magenta) at straight junctions with time (N = 3 repeats). **(b)** Fzd6 and Vangl1 polarized at opposite ends of the cell. Cells co-stained for Fzd6 and Vangl1 (left panel) and the relative ratio of the corresponding proteins along the membrane (right panel) are shown. (N = 6, n = 2130). **(c)** Microtubules polarize along the longest cell axis. To the top, the cells depicting the longest axis (dotted yellow line) and microtubules are color-coded in an angle-dependent fashion. To the right is a scatter plot depicting the angle of the longest cell axis versus the dominant MT angle (n = 10 cells). **(d)** Graphical summary of proposed hypothesis (top) and reductionist numerical model (bottom). Fzd6 (green) and Vangl1 (magenta) are both transported along microtubules from endogenous pools to the surface. Since microtubules align along the longest cell axis and the MTOC is localized in front of the nucleus, we reason that a higher number of MT endings at the cell front yields a bias in the exocytosis rate of core PCP proteins along the front-to-back axis. Once incorporated into the plasma membrane, Fzd6 and Vangl1 augment endocytosis rates of each other (depicted here as lateral inhibition). As Fzd6 is available in higher levels (i.e., retention of Vangl1 through Coronin1c), lateral inhibition combined with polarized transport augments patch formation of Fzd6 at the leading edge. **(e)** The reductionist numerical model recapitulates the polarization of core PCP proteins through biased exocytosis. To the left, the kymograph of localization of Fzd6 (green) and Vangl1 (magenta) over 200 time-steps using a Fzd6:Vangl1 ratio of 3:1 and a 10% bias in exocytosis rate along the flow axis is shown. To the right, an analysis of the Fzd6 vs. Vangl1 ratio along the front-to-back axis is plotted (n = 8 simulations). Reminiscent of live cells, we observe the formation of Fzd6 patches at the leading edge (left) and the polar distribution of Vangl1 vs. Fzd6 (right) is observed. Scale bars, (a,b,c) 10 μm.

In a final set of experiments, we tested the mechanisms by which Coronin1C-dependent changes in the subcellular distribution of Vangl1 may translate to the polarization of PCP proteins within the plasma membrane. A hallmark of core PCP proteins is the formation of two distinct complexes at opposing cellular sites (**Fig. S1**). This mutual exclusion results from endocytosis of the less stable PCP complexes (Cho et al., 2015; Strutt and Strutt, 2008; Strutt et al., 2011) and endosomal microtubule-based trafficking (Matis et al., 2014; Olofsson et al., 2014; Shimada et al., 2006).

Because in endothelial cells, microtubules align along the flow axis (**Fig. 4c** and **Fig. S8b,c**), we reasoned that a cell-shape dependent bias in exocytosis may be sufficient to mediate the formation of mutually exclusive regions of Fzd6 and Vangl1 along the same axis. To probe this hypothesis, we developed a reductionist numerical model (**Fig. 4d, bottom**), in which we assumed the same lifetimes for Vangl1 and Fzd6 at the surface. We further considered that adjacent Vangl1 and Fzd6 mutually reduced their respective lifetimes. To account for lateral protein diffusion, the effect was averaged over 5 neighboring positions. To recreate the conditions of endothelial cells at the onset of flow (**Fig. 2b, middle**), we assumed a 3:1 ratio of Fzd6 vs. Vangl1 and a 10% bias in exocytosis rate along the flow axis. Strikingly, we found polarization of Fzd6 and Vangl1 along the front-back axis in the model that strongly resembles the situation in cells (**Fig. 4b**). The model also shows that a slight bias in the exocytosis distribution, paired with a lateral inhibition between Fzd6 and Vangl1, was sufficient to reconstitute the phenotype observed in the vascular system.

### Vascular remodeling in 3D depends on core PCP proteins Fzd6 and Vangl1

Exposure to shear forces causes individual endothelial cells to elongate and migrate along the flow axis (Levesque and Nerem, 1985; Noria et al., 1999). To aid this process, Fzd6 enriches at the leading edge of migrating endothelial cells, where it exhibits dynamic junctional remodeling to facilitate cell movement. As a consequence of this differential polarization and adhesion, cells travel in streams along the flow axis (**Fig. 5a**). Notably, previous studies established that polarized transcellular stability is also required for sprouting in vivo and in vitro (Bentley et al., 2014). Similar to shear force-dependent cell rearrangements, sprouting further requires cytoskeletal polarization and reorganization of the VE-cadherin–containing complex (Cao et al., 2017). Considering the similarities, we reasoned that Vangl1 and Fzd6 may also contribute to vascular sprouting and angiogenesis.

**Figure 5:**
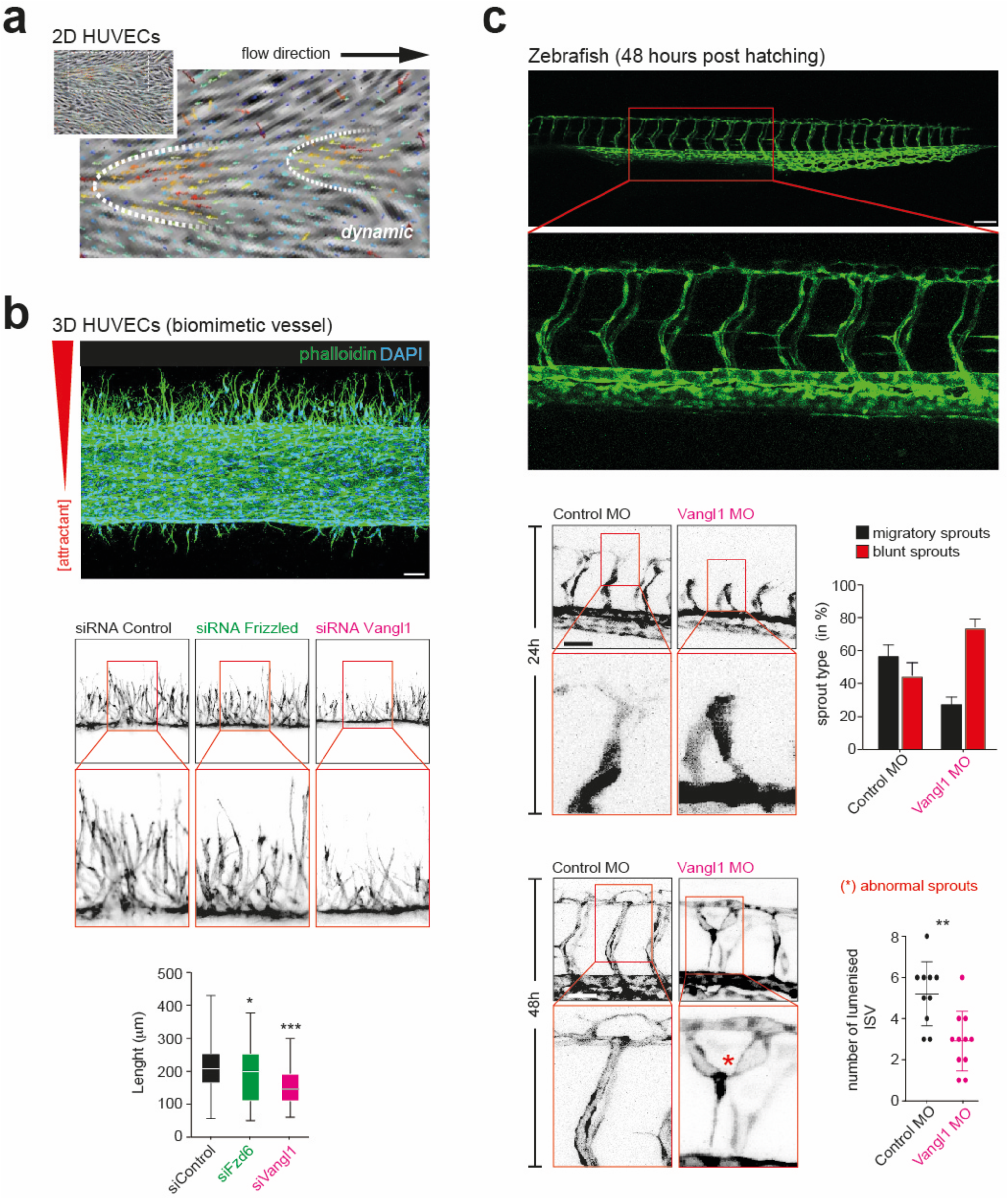
Core PCP proteins regulate endothelial sprouting in vitro and in vivo. **(a)** Upon exposure to shear forces, individual endothelial cells migrate in small cohorts. Analysis of motion pattern of HUVECs cultured in a 2D microfluidic chamber upon exposure to shear forces (18 dyn/cm^2^). Note that individual cells move collectively. **(b)** Loss of Fzd6 and Vangl1 both reduce sprouting in vitro (Kruskal‒Wallis tests from left to right: * p = 0.0257 and **** p < 0.0001, siRNA Control: N = 3 repeats, n = 409 sprouts; siRNA Fzd6: N = 3 repeats, n = 198 sprouts; siRNA Vangl1: N = 3 repeats, n = 178 sprouts). HUVECs cultured in a biomimetic vessel system were exposed to a chemical cocktail that mediates sprouting. Note the changes in sprouting efficiency in cells lacking Fzd6 (green) and Vangl1 (magenta). **(c)** Loss of Vangl1 reduces vein sprouting in vivo. *vangl1* morpholinos were injected into one-cell stage zebrafish embryos, and tail vein formation was analyzed. Loss of Vangl1 showed a significant increase in blunt sprouts at 24 hpf (middle panels, red). (Two-Way ANOVA: ** p = 0.0077, control morpholinos: N = 3 biological replicates, n = 409 sprouts; *vangl1* morpholinos: N = 3 biological replicates, n = 178 sprouts). At 48 hpf, embryos lacking Vangl1 displayed sprouts with incorrect connections (abnormal sprouts, red), frequently lacking a visible lumen (bottom panel, magenta). (t-test: p = 0.0024, Control morpholinos: N = 3 repeats, n = 10 embryos; Vangl1 morpholinos: N = 3 repeats, n = 11embryos). Scale bar, (b, c, top) 100 μm, (c, middle and bottom) 50 μm.

To validate this hypothesis, we took advantage of a biomimetic 3D system, where HUVECs sprout in a polarized fashion into a gel towards a chemo-attractant (Liu et al., 2021). Consistent with the role of core PCP proteins in this process, we find reduced sprouting and the formation of single-cell clones in cells lacking Fzd6 and Vangl1 (**Fig. 5b**). We then aimed to determine whether the observed role of core PCP proteins in the biomimetic system can also be found in vivo (**Fig. 5c**). We analyzed the zebrafish intersegmental vessels (ISV) development, a bona fide model system of vascular sprouting formation. We focused on the function of Vangl1 due to its strong *in vitro* phenotype and displaying all PCP protein characteristics. A validated antisense morpholino was previously published (Borovina et al., 2010). *vangl1* antisense morpholinos (*MO-vangl1*) and control morpholinos (*MO-control*) were injected into *Tg(kdrl:EGFP)* embryos, a pan-endothelial transgenic line with fluorescent labeling of all endothelial cells (Beis et al., 2005). At 24 hours post fertilization (hpf), embryos injected with *MO-vangl1* displayed a significant increase in shorter blunt sprouts compared to zebrafish injected with *MO-control* (**Fig. 5c, middle**). Sprout morphology showed significant defects in *MO-vangl1* embryos at 48 hpf, frequently forming three-way connections that did not resolve and ISVs with abnormal diameter (**Fig. 5c, bottom** and **Fig. S10a**), thus establishing a dual role of core PCP proteins in the vascular system (i.e., 2D cellular alignment and 3D sprouting).

## DISCUSSION

To accurately adapt to changing conditions, the vascular system continuously senses and adjusts its mechanical properties to physical stimuli from the environment (Aitken et al., 2023; Fang et al., 2019). Naturally, the response of the vasculature varies based on the type of stimuli, yielding a rich repertoire of possible cell states. While functionally and mechanically different, most states share the ability to polarize the cell along the direction of the blood flow. Here, we aimed to investigate how this polarization is accomplished. We find that core PCP signaling promotes cell shape changes and migration dynamics along the flow axis as well as vascular sprouting. Conceptually, our findings yield four main advancements.

First, we introduce flow-dependent mechanical forces as a global cue for the polarization of core PCP proteins in the vasculature. In other multi-cellular systems, multiple, potentially overlapping, actuators have been identified that influence core PCP alignment. These include, among others, expression gradients of Fj and Ds (Galic and Matis, 2015; Matis and Axelrod, 2013), Wnt ligand gradients (Gao et al., 2011; Gros et al., 2009) and mechanical cues (Chien et al., 2015; Guirao et al., 2010; Mitchell et al., 2009; Valet et al., 2022). Complementing these chemical and mechanical cues, we demonstrate shear force as a global cue to polarize core PCP proteins and induce changes in cell shape and migration. Since initial cell elongation precedes polarization and surface exposure of Vangl1, core PCP proteins augment an initial polarization along the flow axis. What induces this initial elongation remains elusive.

Second, our results present a novel mechanism for the symmetry-breaking of core PCP within a cell. In epithelial cells, PCP proteins are initially uniformly distributed but become polarized before morphological polarization. Fz, Dsh and Dgo become highly enriched at one side of the cell cortex, while Vangl and Pk localize to the opposing apical side of the cell (Bastock et al., 2003; Tree et al., 2002). Remarkably, in HUVECs, Vangl1 and Fzd6 protein dynamics significantly deviate from this pattern. Unlike in other cellular systems, surface delivery of individual core PCP proteins in the endothelium is sequential. The numerical model indicates that biased exocytosis upon flow-dependent alignment of MTs is sufficient for a polarized enrichment of proteins. To our knowledge, this is the first time that the frequency of core PCP protein incorporation has been introduced as the source of symmetry breaking. Notably, junctional incorporation of Vangl1 is not only concomitant with the polarization of Fzd6 but also coincides with a reduction of the total levels Fzd6 (**Fig. 3a**). While this study introduces Coronin1C as a mediator for the delayed recruitment of Vangl1 to the cell surface, we do not rule out additional signaling circuits influencing the equilibrium between endo- and exocytosis or the surface retention of Vangl1 and other core PCP proteins.

Third, our data suggests a dual role for core PCP signaling within the vasculature. In the planar vascular system, the loss of core PCP proteins delays and weakens cell elongation and migration along the cell axis. Considering that Fz/Dvl promotes actin polymerization while Vangl/Pk antagonizes it (Davey and Moens, 2017) and **Fig. S9**), we propose that the observed enrichment of Fzd6 and Vangl1 at opposite sites augments polarized cell dynamics along the flow axis. Importantly, we also observe disrupted sprouting upon loss of Vangl1. Impairment in 2D alignment and 3D sprouting not only indicate a shared mechanism governing both processes but also raise the possibility of interdependence between the two phenomena.

Finally, loss of Vangl1 is associated with an increase in pro-inflammatory gene expression (**Fig. S10b**), suggesting elevated stress levels in cells incapable of aligning along the flow axis. Physiologically, it raises the question of whether core PCP signaling may participate in pathophysiological conditions where blood flow has been impaired. Previous studies established that long-term unidirectional flow within the physiological range promotes quiescence with low proliferation and inflammatory gene expression (Chakraborty et al., 2012; Wang et al., 2013). Conversely, low flow or changes in the flow direction (multidirectional or oscillatory flow) promote proliferation and inflammation (Bondareva et al., 2019; Chakraborty et al., 2012; Wang et al., 2013). It is thus plausible to envision impaired core PCP signaling in hyperlipidemia, smoking, and diabetes, where flow patterns have gone awry.

## Acknowledgments

We would like to thank the members from the Matis, Galic, Raz, and Klingauf labs for their critical feedback and insightful discussions. Verena Stegemann is thanked for help with experiments in the biomimetic 3D system. This work was supported by funds from the DFG to FG (CRC1348/B12; SF1450/N04), BT (CRC1348/A07), HS (DFG grants SCHN 430/6-2 and SCHN 430/9-1), MG (GA 2268/3-1; GA 2268/4-1; CRC1348/A06), and MM (MA 6726/3-1, EXC-1003/FF-2015-07), from the DZHK to HG (81X3100105) and MOS (DZHK, 81×2100151), as well as by funding from the Medical Faculty of the University of Münster to FG (IMF IGU-122208), MG (IMF IGA-121610) and MM (IMF IMA112016 and IZKF Mat1/027/21).

## Author contributions

LG performed all experimental work unless stated otherwise. TS developed image analysis software with input from MM and MG. LG, JL, MOS, and HS performed 2D culture experiments. EBA and HG performed VE-Cadherin proximity ligation experiments, and OP and PM performed the proximity ligation MS experiment and analysis. LG and BM performed Vangl1-specific MS analysis. LG and BT performed 3D culture experiments. LG, MM and FG performed zebrafish experiments. MM and MG developed and implemented the numerical model. LG, MM and MG designed experiments, discussed results and wrote the manuscript with input from all authors.

## MATERIAL AND METHODS

### Cell culture

For all flow experiments with the BioTechFlow system (MOS Technologies, Telgte, Germany), HUVECs from passage #1 were seeded on crosslinked gelatin-coated culture dishes as previously described (Taha et al., 2019). The use of HUVEC was according to the principles outlined in the Declaration of Helsinki and was approved by the ethics boards of the WW-University of Muenster (2009-537-f-S). HUVECs were seeded at 4×10^4^ cells/cm^2^ to achieve confluency (1-1,2 x10^5^ cells/cm^2^) 3 to 4 days later. Brightfield images of fluid shear stress experiments were acquired using the BioTechFlow (BTF) system equipped with a modified inverse Zeiss Axiovert 20 microscope. Shear stress was applied in endothelial cell growth medium (PromoCell, C-22010) supplemented with 3% polyvinylpyrrolidone (PVP, Sigma-Aldrich-Aldrich, 9003-39-8) in the presence of 5% CO_2_. Experiments and corresponding controls were always performed in parallel.

For all flow experiments with the Ibidi Pump System (Ibidi, 10902) (Experiments in Fig. 3b,c,d,g), HUVECs were purchased as cryo-conserved pools (Promocell, C-12208). Cells were cultured in an incubator at 37°C and 5% CO_2_ with 100% humidity in a 1:1 mixture of EGM-2 with M199 (Thermo Fischer, 31150022) supplemented with 10% fetal calf serum (FCS; Sigma-Aldrich, F7524-500ML), 20 µg/ml gentamicin (Thermo Fischer, 15710049), 15 µg/ml amphotericin B (Thermo Fischer, 15290026), and 100 IE heparin (Sigma-Aldrich, H3149). For flow experiments, Endothelial Growth Medium 2 (EGM-2; Promocell, C-22011), supplemented with EGM-2 serum, was used. Cells were used at passage 2-4 in all experiments. Again, experiments and corresponding controls were always performed in parallel.

### RNA isolation and quantitative RT-PCR

Total RNA was isolated using the RNeasy mini kit (Qiagen, 74104), and cDNA was synthesized with oligo(dt) primers and Superscript-III-RT (Invitrogen, 18080093). Expression of mRNA was quantitatively assessed by real-time PCR using KAPA SYBR FAST (Sigma-Aldrich, KK4608) in a Biorad iCycler IQ (Biorad, 582BR). Primer sequences can be found in **Table S1**.

### siRNA transfection

The transfection of siRNA constructs (**Table S2**) was performed by AMAXA transfection according to manufacturing protocol (Lonza, VPB-1002). Briefly, 10^6^ suspended HUVECs were diluted in 100 μL transfection buffer containing 20 pmol (0.26 μg) siRNA. After electroporation, cells were seeded at subconfluent densities and incubated for 24 or 48 hours in the EGM-2/M199 medium. For shear stress conditions, HUVECs were transfected by magnet-assisted transfection (MATra; Promocell, CT-2021-020) according to the manufacturer’s protocol. In short, 3 μg siRNA was mixed with 9 μL MATra siRNA transfection solution and added to 2 ml EGM2 medium. This mixture was added to confluent HUVECs grown on a gelatin coated dish. The dish was placed on the supplied magnet for 20 min at 37°C. Subsequently, the medium with MATra and siRNA solution was removed, and a complete Promocell medium was applied. Shear stress conditions were introduced 24 hours after MATra transfection.

### Plasmid transfection

For overexpression of Vangl1 (OriGene Technologies, RC209024), Lipofectamine 2000® (Thermo Fischer, 11668030) was used. One hour before transfection, the medium was replaced with EGM-2 basal medium supplemented with 2% FCS. Two separate dilutions were prepared: 125 ng of plasmid in 16.67 μL of Optimem® Reduced Serum Medium (Thermo Fischer, 31985062) and 0.115 μL of Lipofectamine® in 16.67 μL of Optimem®. After 5 minutes at 25°C, the diluted DNA was added to the diluted Lipofectamine® and incubated for 20 minutes at 25°C. Then, 66.67 μL of EGM-2 + 2% FCS was added to the DNA-lipid complexes. The mixture was applied to confluent HUVECs grown on a gelatin-coated μ-Slide. After 3 hours at 37°C, the medium containing the DNA and Lipofectamine solution was removed and replaced with the complete EGM-2 medium. Shear stress conditions were introduced 24 hours post-transfection with Lipofectamine.

### Fixation protocols

For immunocytochemistry, cells were washed shortly with DPBS, fixed in 2% PFA (Roth, 4235.1) for 10 min at RT, permeabilized with 0.0.5% Triton-X100 (Sigma-Aldrich, 93443) in PBS for 10 min at RT, followed by a short washing step and blocked 1 hour at RT in 5% serum, 0.1% Tween (Applichem, A4974,0250), 300 mM Glycine (Applichem, A1067.5000) in PBS. Cells were incubated with primary antibodies overnight at 4°C, washed with DPBS 3 times and incubated with relevant fluorescent-conjugated secondary antibodies (**Table S3**). After Dynamin treatment (Fig. 3c), cells were fixed with 4% PFA in PBS at room temperature for 10 minutes and washed 3 times with PBS. They are blocked with 5% serum (Abcam, ab7481) and 0.05% saponin (Sigma-Aldrich, S7900-25G) in PBS for 1 hour at RT and again washed 3 times with PBS. Primary antibodies were diluted in PBS containing 0.05% Saponin and 1% serum and left overnight at 4°C. The next day, cells were washed 3 times with PBS and incubated with corresponding secondary antibodies in 0.05% saponin and 1% serum. The next day, the cells were washed with DPBS 3 times and incubated with relevant fluorescent-conjugated secondary antibodies.

### Microscopy and image analysis

The stained cells were mounted with Ibidi mounting medium (Ibidi, 50001) and imaged with a Zeiss 710 Microscope (Carl Zeiss) and a Plan-Neofluar 63x (NA 1.4) oil immersion objective. Images were collected using ZEN software. All images were processed using Fiji/ImageJ (Version 1.49p) (Schindelin et al., 2012) or Matlab (2019b). Ratiometric analysis, velocity maps, track analysis, and vector analysis were done in Matlab using custom-made codes. All source code will be made publicly available on our homepage (https://campus.uni-muenster.de/en/impb/das-institut/nanoscale-forces-in-cells/software/) upon publication.

### Protein extraction and Western blot

For immunoblotting, cells were lysed in RIPA buffer (Thermo Fischer, 89900) containing protease inhibitor cocktail (Thermo Fischer, 78430), phosphatase inhibitor (Thermo Fischer, 78420), and 100 mM PMSF (Sigma-Aldrich, 329-98-6). Protein concentrations were measured using the Pierce BCA Protein Assay Kit (Thermo Fischer, A55864). Samples were further diluted with SDS-loading buffer, and SDS-PAGE was performed using 8–12% Bis-Tris Protein Gels. Proteins were transferred to PVDF membranes (Merck, IPFL00010) by semi-dry blotting. Membranes were blocked with Intercept® (TBS) Blocking Buffer (Li-cor, 927-60001) for 1 hour at room temperature (RT) and primary antibodies were incubated overnight at 4°C in a 1:1 mixture of blocking buffer and TBS with 0.1% Tween (Applichem, A4974,0250). Host matching fluorescent secondary antibodies IRDye 800CW (LiCor; Donkey anti-mouse 800CW, 926-32212; Donkey anti-goat 800CW, 926-32214; Donkey anti-rabbit 800CW, 926-32213) and IRDye 680RD (Li-cor, Donkey anti-goat 680RD, 926-68074; Donkey anti mouse 680RD, 926-68072; Donkey anti rabbit 680RD, 926-68073) were diluted 1:10,000 in TBS-T and incubated 1 hour at RT. Membranes were imaged with a Li-cor imaging system (LI-COR Biosciences, Odyssey Infrared Imaging System Application Software Version 3.0)

### Cell fractionation by sucrose density gradient centrifugation

For subcellular fractionation, HUVECs were washed one time with ice-cold PBS pH 7.4 and scraped into 1 ml of ice-cold sucrose buffer (0.25 M Sucrose, 10 mM Tris-HCl pH 7.4, 1 mM magnesium acetate and mixture of protease/phosphatase inhibitors). The cells were homogenized by 6 passages through a 25G needle. Homogenates were centrifuged at 2500 x g for 15 min at 4°C and supernatants were loaded on sucrose step gradients consisting of (from top to bottom) 0.25 M (0.5 ml), 0.5 M (1.6 ml), 0.8 M (1.6 ml), 1.16 M (2 ml), 1.3 M (2 ml), and 2 M (1.2 ml) sucrose in 10 mM Tris-HCl (pH 7.4) and 1 mM magnesium acetate. The gradients were centrifuged at 39,000 rpm for 2.5 h at 4°C using a SW41Ti rotor (Beckman Instruments), and 21 fractions of 0.5 ml were collected. Samples were mixed with 5X loading buffer and analyzed by SDS-PAGE and Western blot. For mass spectrometry analysis, 350 μL of Vangl1 positive fractions were diluted with 10 volumes of Tris buffer and pelleted in an ultracentrifuge equipped with a SW41Ti rotor (Beckman Instruments) at 25.000 rpm for 1 hour at 4°C. The pellet was resuspended in 50 μL 1x loading buffer.

### TurboID proximity labeling with VE-Cadherin as bait

Adeno-associated viruses for VE-Cadherin proximity labeling experiments were generated using the ViraPower Adenoviral Expression System (Invitrogen, K493000). Briefly, the previously generated pAd::VE-Cad::TurboID plasmid using TurboID plasmid backbone (Branon et al., 2018) was digested via PacI restriction, and the linearized DNA was transfected into 293A cells (Invitrogen, R70507). Crude viral extract from transfected cells was harvested a day later, and the transfection step was repeated with the cell lysate to amplify the viral concentration titer. Following the second harvest and titer measurement, the amplified adenovirus stock was aliquoted and stored at −20C for long-term storage. HUVECs, seeded into a 6-well plate and incubated in EGM2 media until 80% confluency, were transduced with adenoviruses overnight and were then washed 3 times with EGM2 media before further processing for experiments to remove the residual AAVs.

Following pAd::VE-Cad::TurboID adenoviral transduction, HUVECs were seeded into ibidi perfusion slides for 0h static and 24h flow conditions as previously described (Vion et al., 2021), at a concentration of 2,5*105 cells per slide, while Ibidi 2-well slides were used for 24h static condition, at 4*105 cells per well. 4 µ-Slides (Ibidi, 80176) or 2 2-well plates (Ibidi, 80286) were seeded per condition. For flow conditions, perfusion flow units were set up as previously described, and cells were perfused at 20 dyne/cm2. 2 hours before the condition timepoint underflow, flow pumps were briefly stopped, and stock biotin was directly added on the top of the syringes inside the incubator to bring the media to a final biotin concentration of 500 µM, followed by perfusion until the desired timepoint. For static samples, biotin was diluted in warm EGM2 media to a final concentration of 50 µM, prior media was replaced with biotinylated media, and samples were similarly incubated for 2 hours.

For protein extraction and subsequent biotinylated protein immunoprecipitation, a modified protocol based on Branon et al. was established (Branon et al., 2018). Adherent HUVECs were washed 1x with warm PBS, followed by addition of 200 µL modified RIPA buffer(50 mM Tris-HCl (pH 7.2), 150 mM, NaCl, 1% NP-40, 1 mM EDTA, 1 mM EGTA, 0.1% SDS, 1% sodium deoxycholate, 1:500 Benzoase (Millipore, 70746), Halt Complete protein inhibitor (Thermo Scientific, 78429)) per sample and harvesting by rapid pipetting of ibidi flow slides with a 1 mL syringe with Luer-Lok (Fischer scientific, 309628), or with a cell scraper for 2 well plates. Streptavidin-tagged agarose beads (Thermo Scientific) were used as an alternative to magnetic beads used on the original protocol. Prior to pulldown, the beads were prepared by washing them twice in modified RIPA buffer, removing liquid with a Hamilton SampleLock syringe (Hamilton, 2.5 mL blunt tip, 22 gauge, 20353), and resuspending again in RIPA buffer (50% v/v). The subsequent wash, bead pulldown, digestion, and elution steps followed the original protocol steps, using Hamilton pipetting to discard supernatant during wash steps in order to keep bead loss to a minimum.

Tryptic on-bead digestion was carried out following the protocol from Hubner et al. (Hubner et al., 2010). The peptides were then desalted on stage tips as described by Rappsilber et al. (Rappsilber et al., 2007). Liquid chromatography-mass spectrometry (LC-MS) measurements were conducted using an Exploris 480 mass spectrometer (Thermo Fisher Scientific) connected to an EASY-nLC 1200 system (Thermo Fisher Scientific). A 110-minute gradient was applied, and the mass spectrometer operated in data-dependent mode.

For the analysis, MaxQuant version 2.0.3.0 (Cox and Mann, 2008) was used, employing iBAQ-based quantitation and utilizing the match-between runs algorithm. Carbamidomethylation was set as a fixed modification, and oxidized methionine was set as a variable modification. The Andromeda search engine was used with a Uniprot human database (2018-07) combined with common contaminants. Proteins were filtered to include only those with at least two peptide identifications across all runs. Log2 iBAQ values were used for quantitation. A valid value filter required at least 50% values across all samples and outlier replicates have been removed. Intensities were median-non-zero normalized. The remaining missing values were imputed using down-shift imputation. Downstream analysis was performed in R using two-sample moderated t-statistics with the limma package (Ritchie et al., 2015).

### Inhibition of endocytosis by Dynasore

HUVECs from passage #1 were seeded on crosslinked gelatin-coated Ibidi slides in EMG2 medium. Cells were grown for 4 days, with the medium changed every 2 days. To block endocytosis, a complete EMG2 medium was supplemented with 40 μM Dynasore or an equivalent concentration of only DMSO and added to the cells. The cells were incubated for 4 hours in an incubator at 37°C and 5% CO_2_. Subsequently, the cells were fixed with 4% PFA and processed as described in the fixation protocol.

### Numerical model

The cell periphery was considered a chain of equidistant positions. The simulation was initiated by randomly picking positions and randomly depositing one particle (either Vangl1 in magenta or Fzd6 in green). Each particle was allocated a random lifetime. All particles with expired lifetime were removed and randomly placed back in a free position. Upon arrival, the new lifetime was determined depending on the number of same-colored particles in the vicinity: if less than 4 particles were in the vicinity, the particle was annotated with a shorter lifetime. The simulation was left to evolve for 200 timesteps. The source code will be made publicly available on our homepage (https://campus.uni-muenster.de/en/impb/das-institut/nanoscale-forces-in-cells/software/) upon publication.

### Ex vivo angiogenesis

Microfluidic devices mimicking sprouting angiogenesis in vitro were prepared as previously described (Nguyen et al., 2013). In short, the device consisted of two patterned layers of PDMS, which were molded from photographically patterned silicon masters and sealed against a glass coverslip. The PDMS and glass surfaces were functionalized with an aqueous 0.1% (w/v) poly-L-lysine and, subsequently, 1% (w/v) glutaraldehyde solution. To form tubular channels, needles of 400 μm diameter were coated with a 0.4% aqueous solution of BSA, sterilized using UV light and inserted into the device. A 2.5 mg/ml collagen solution was cast inside the device and allowed to polymerize for 30 min at 37°C. The resulting gels were hydrated in PBS overnight and washed thoroughly with PBS before needle extraction. A solution of 10^6^ cells/ml medium was added to a reservoir, and cells were allowed to attach to the bottom side of the channel for 10 min, followed by seeding of the top channel side for another 10 min. After washing with media, the devices were placed on a platform rocker to initiate gravity-driven flow. 5 hours after seeding, a growth factor cocktail consisting of 75 ng/ml VEGF (R&D Systems, VE-293), 10 ng/ml PMA (Sigma-Aldrich, P1585), and 500 nM S1P (Cayman Chemical) in media was introduced to the second channel to induce angiogenic sprouting. Media and cocktail were exchanged daily, until cells were fixed in 4% PFA after 2 days of culture. For imaging, fixed sprouts were first permeabilized in 0.1% Triton-X100 in PBS for 1 hour and then stained with Phalloidin-635Star (Abberior, ST635-0100-20UG) overnight in PBS with 0.1% Tween at 4°C.

### In vivo angiogenesis

To fluorescently label endothelial vascular beds, the pan-endothelial transgenic line *Tg(kdrl:EGFP)^s843^* was used (Beis et al., 2005). Embryos were raised in 0.3× Danieau’s solution (17.4 mM NaCl, 0.21 mM KCl, 0.18 mM Ca(NO3)2, 0.12 mM MgSO4·7H2O) at 28°C. The zebrafish were handled according to the regulations of the state of North Rhine-Westphalia, supervised by the veterinarian office of the city of Muenster. Control morpholino (5’-CCTCTTACCTCAGTTACAATTTATA-3’), and vangl1 morpholino (5’-CATGGCAATGGCGTCTGTGCTG-3’) were injected at a concentration of 300 µM, as previously published (Borovina et al., 2010). Per zebrafish, 1 nL of morpholino-containing solution was injected into the yolk of one-cell stage embryos.

### Statistical analysis

All the datasets presented in this work were first tested for normal distribution using D’Agostino & Pearson or Shapiro-Wilk normality tests. Statistical significance was determined using two-tailed unpaired/Student’s t-test or ordinary one-way ANOVA (with Tukey’s multiple comparisons) when the data were distributed normally for two or more than two groups, respectively. Two-tailed Mann-Whitney U test or Kruskal-Wallis test (with Dunn’s multiple comparisons) were performed when datasets were not distributed normally. All the graphs with scatter dot plots show mean values with red lines. Boxes in all box plots extend from the 25th to 75th percentiles, with a line at the median. Whiskers show min and max values. All the bar graphs indicate mean ± sd. The significance levels in all the graphs are as follows: n.s. (nonsignificant), * (p ≤ 0.05), ** (p ≤ 0.01), *** (p ≤ 0.001) and **** (p ≤ 0.0001). The statistical tests were performed using Prism7 (version 7.0d for Mac OS X, GraphPad Software). All experiments presented in the manuscript were repeated at least in three independent experiments/biological replicates. The experiments were not randomized, and the sample size was not predetermined.

## Data availability

The data that support the findings of this study are available from the corresponding authors upon reasonable request. No restrictions apply.

## Competing interests

The authors declare no competing financial interests.

## Supplementary Information

### SUPPLEMENTARY FIGURE LEGENDS

**Figure S1:**
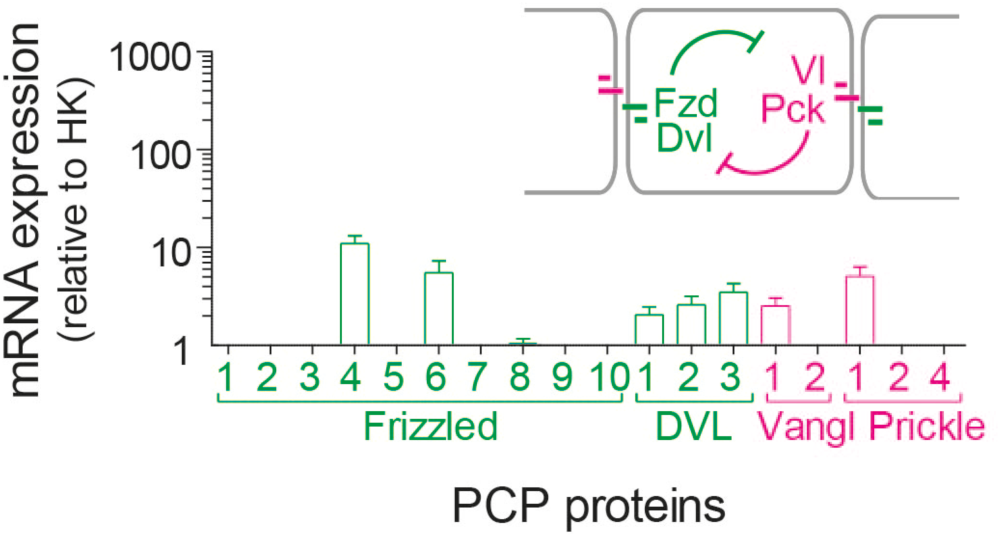
Core PCP proteins are expressed in HUVECs. Real-time PCR shows high gene expression levels for transcripts of Fzd6, Dvl1-3, Vangl1 and Pk1.

**Figure S2:**
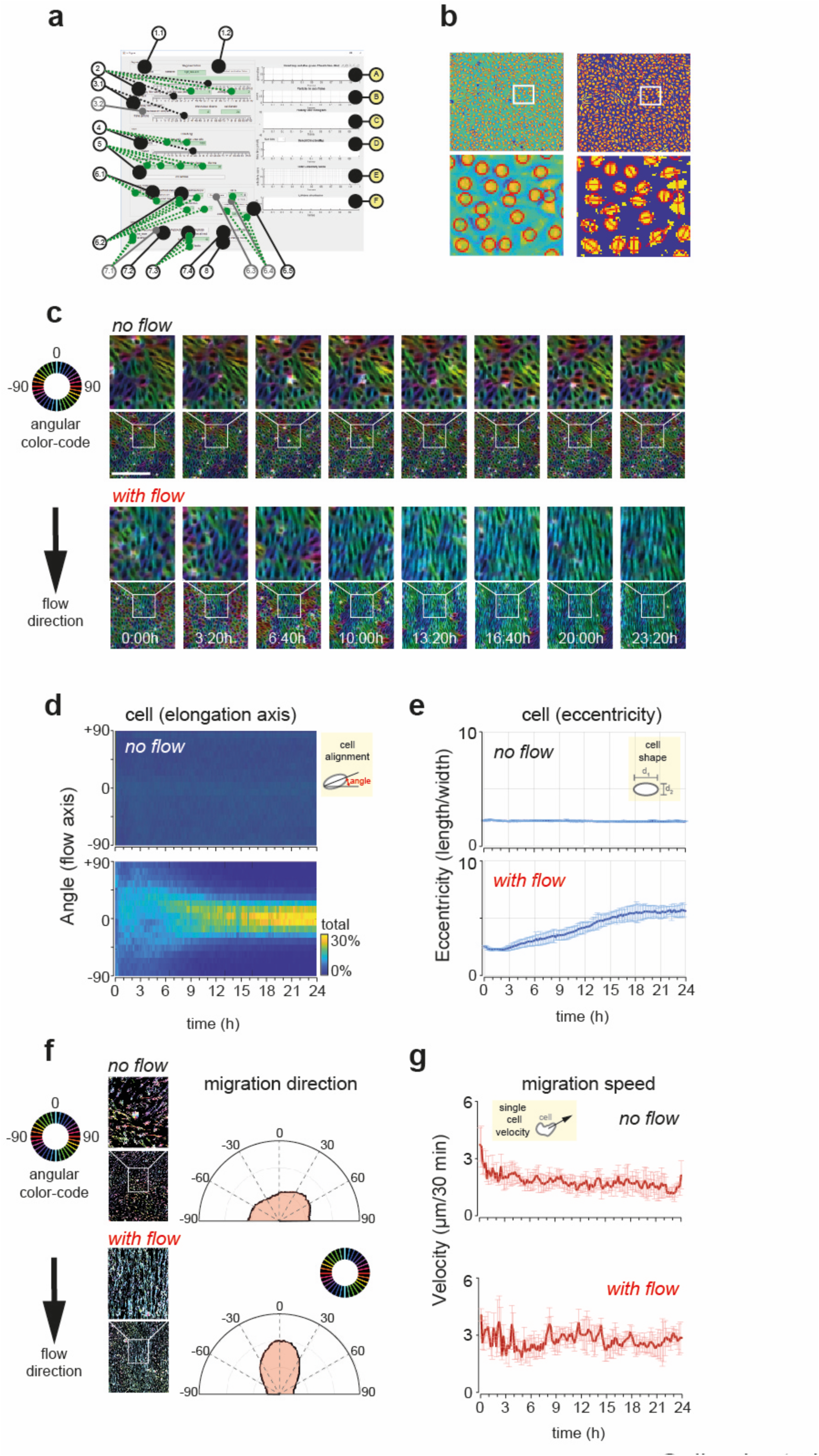
Custom-made image analysis software for the analysis of collective cell dynamics. **(a)** Overview of the graphical interface, which allows the control of individual aspects of cell tracks. For details on handling, please refer to supplemental material. (**b)** Automatic detection and alignment of principal axis from grey-scale brightfield images. **(c)** Fluid shear forces cause cell elongation. Cells in the absence (top) and presence (bottom) of fluid shear forces are shown over time. **(d)** Angular distribution of longest axis over time. As before, cells in the absence (top) and presence (bottom) of fluid shear forces are shown. Note the increase in cell alignment along the flow axis over time. **(e)** Cell elongation, the ratio of the longest vs. shortest perpendicular diameters, was monitored and quantified over 24 hours. No changes are seen for cells in the absence of flow (top panel, N = 3 biological repeats, n = 872 cells). In contrast, cells exposed to fluid shear forces of 18 dyn/cm^2^ displayed a monotonous elongation that started 3 hours after flow induction and plateaued after 18 hours (bottom panel, N = 3 biological repeats, n = 906 cells). **(f)** Direction of cell migration in the absence (top) and presence (bottom) of flow. **(g)** Quantification of migration speed. Single-cell speed of HUVECs in the absence of flow (top panel, N = 3 biological repeats, n = 872 cells) and the presence of fluid shear forces (bottom panel, N = 3 biological repeats, n = 906 cells) are shown. Scale bar, (**c**) 100 μm.

**Fig. S3:**
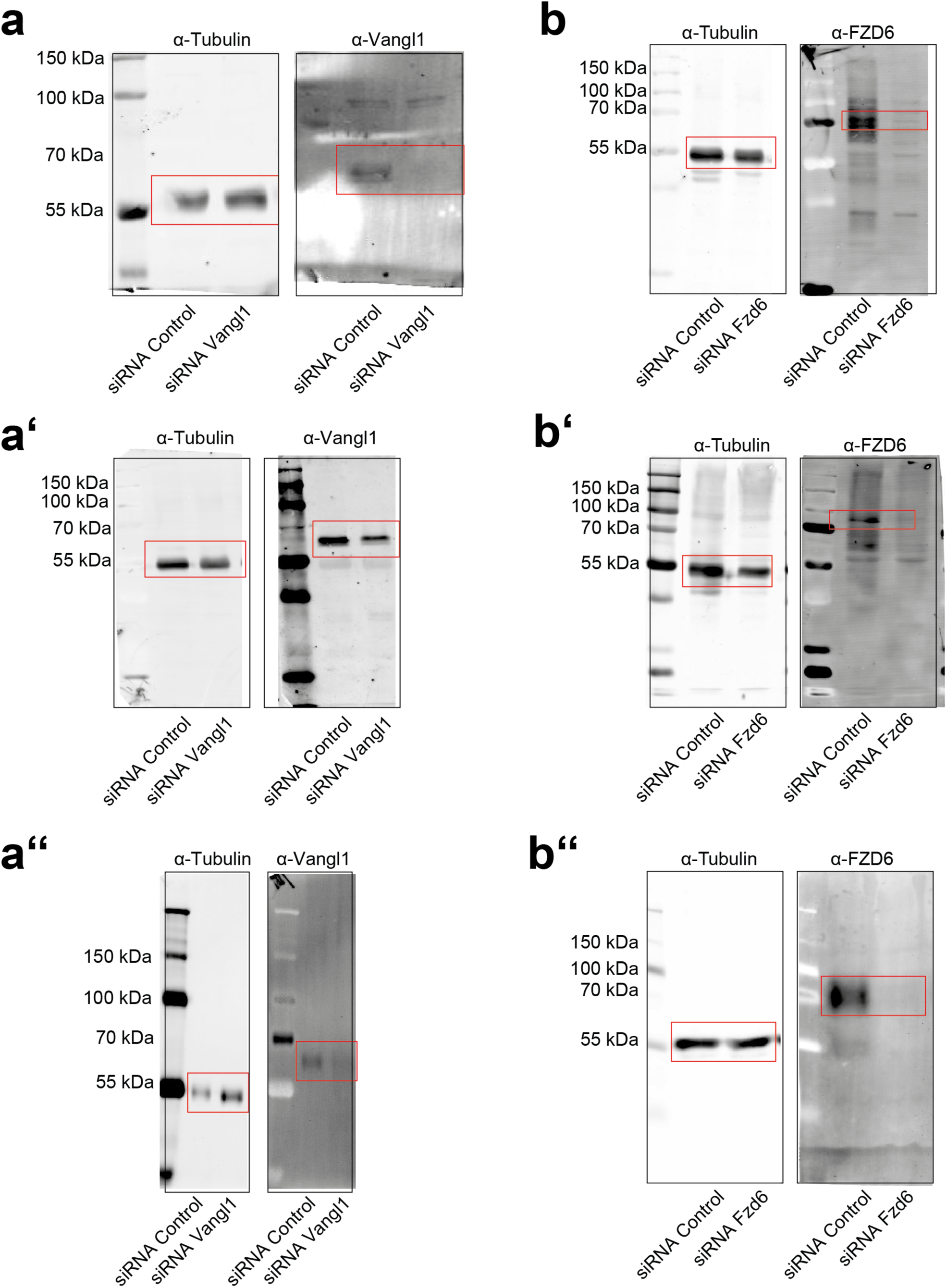
Validation of siRNA knockdown efficiency. **(a)** Western blot of HUVECs transfected with siRNA directed against Control (left) and Vangl1 (right) show a reduction in Vangl1 protein levels. Whole membranes stained with antibodies directed against Tubulin and Vangl1 are shown from left to right. From top to bottom, membranes for three independent biological repeats are shown. **(b)** Western blot of HUVECs transfected with siRNA directed against Control (left) and Fzd6 (right) show a reduction in Fzd6 protein levels. Whole membranes stained with antibodies directed against Tubulin and Fzd6 are shown from left to right. From top to bottom, membranes for three independent biological repeats are shown.

**Figure S4:**
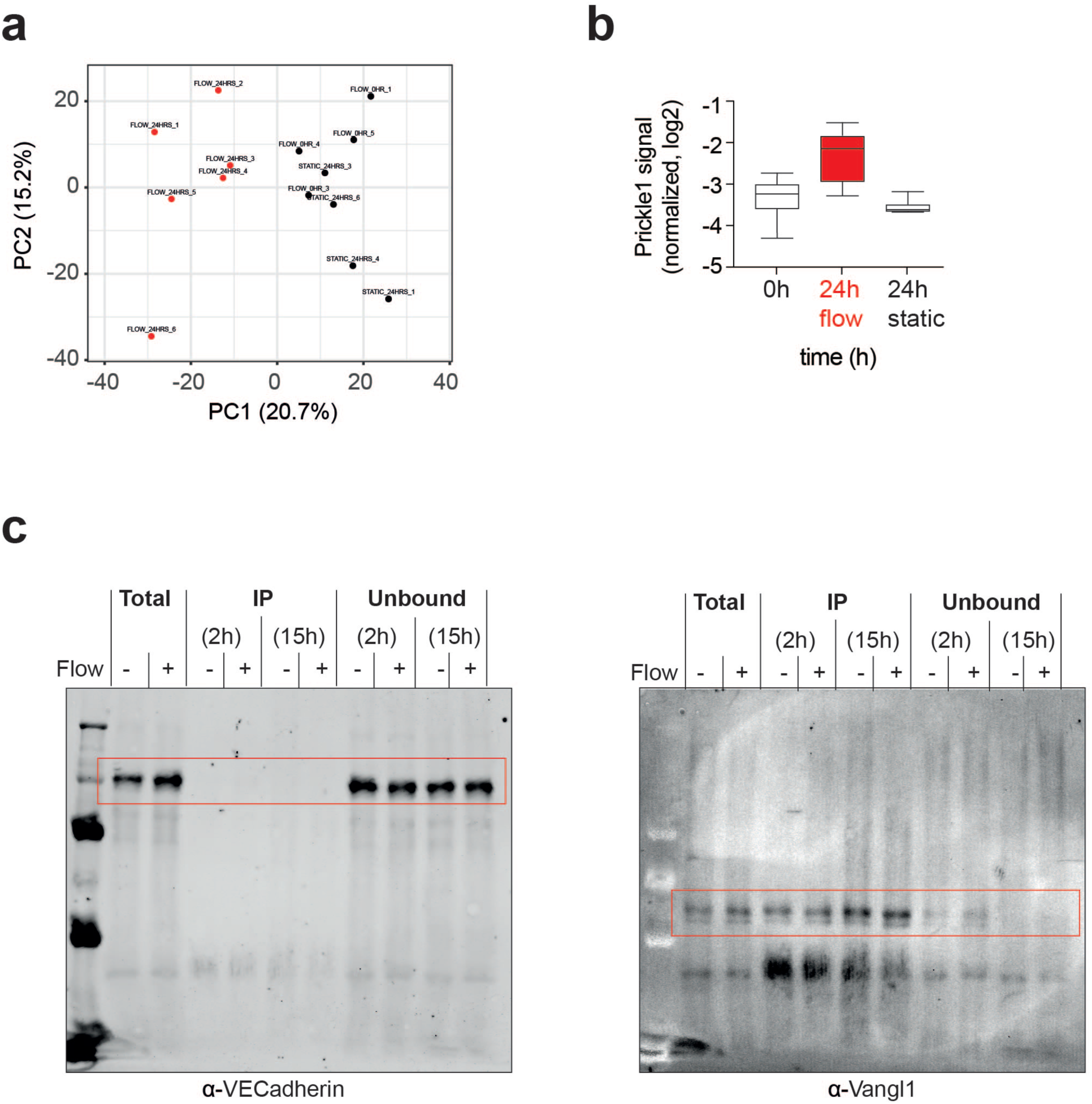
Interaction studies of VE-Cadherin and Vangl1 in the presence and absence of flow. **(a)** Principal component analysis of proximity ligation, using VE-Cadherin as bait in the presence and absence of flow. **(b)** For Prickle1, we find enrichment at the cell surface only upon flow induction (0h, N = 4 repeats; 24h flow, N = 6 repeats; 24h static, N = 4 repeats). **(c)** Vang1 and VE-Cadherin do not interact with each other. Proximity ligation uses VE-Cadherin as bait in the presence and absence of flow. Total lysates were cultured for 2 hours (i.e., 2h) or 15 hours (i.e., 15h) with magnetic beads conjugated to Vangl1 as bait and then subjected to pulldown. Whole membranes treated with antibodies directed against VE-Cadherin (left) and Vangl1 (right) are shown. In each membrane, from left to right, total lysate (± flow), IP for 2 and 15 hours (each ± flow), and flow-through for 2 and 15 hours (each ± flow) are depicted.

**Figure S5:**
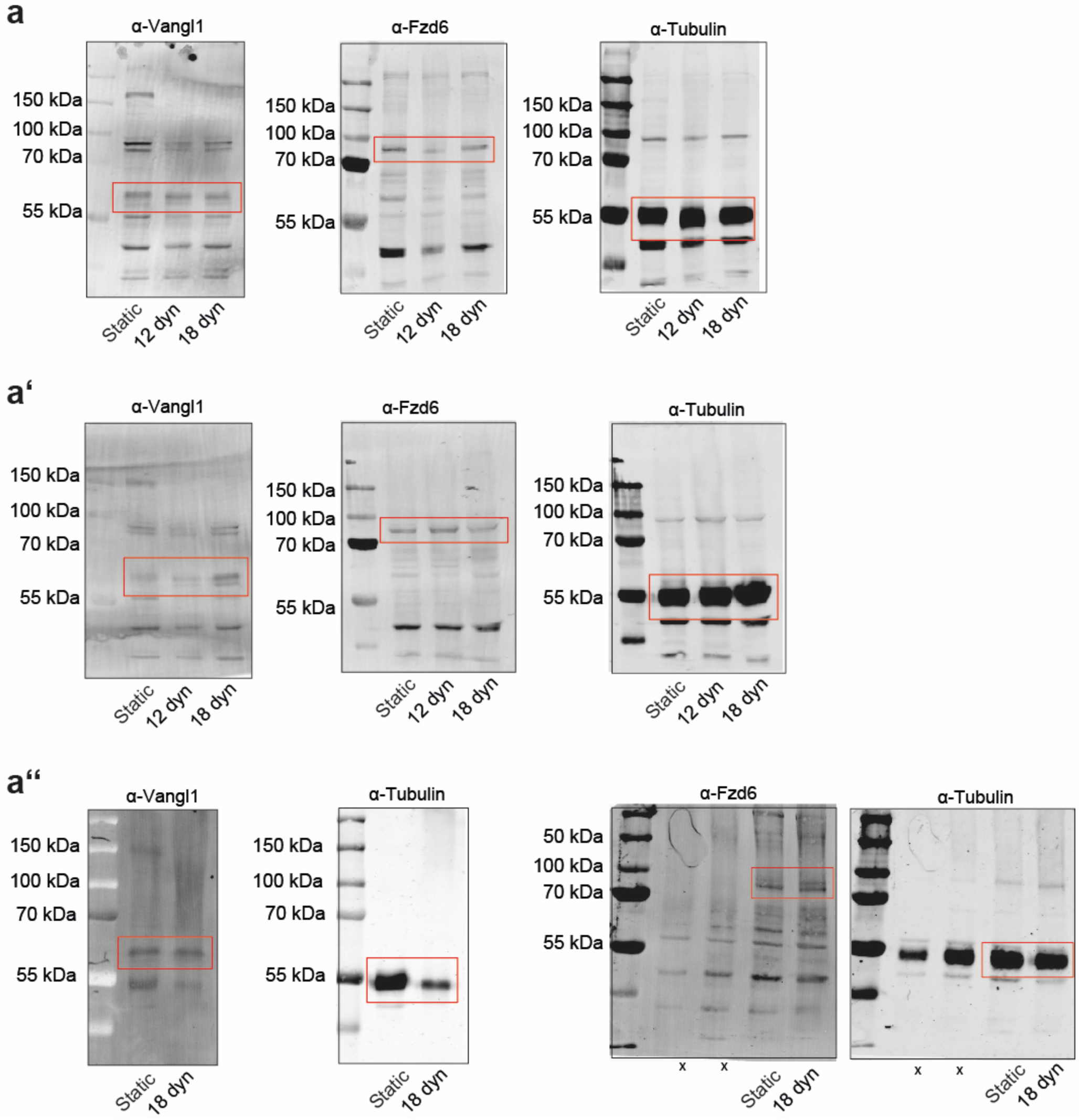
Validation of core PCP protein levels in the presence and absence of shear forces. Western blot depicting HUVECs cultured in the absence of fluid shear forces (left line), exposed to 12 dyn/cm^2^ (middle line), or for 24 hours to 18 dyn/cm^2^ (right line). Total lysates were loaded on a gel and stained with antibodies against Vangl1 (left blot), Fzd6 (middle blot) and Tubulin (right blot). From top to bottom, membranes for three independent biological repeats are shown (a-a’’).

**Supplemental Figure S6:**
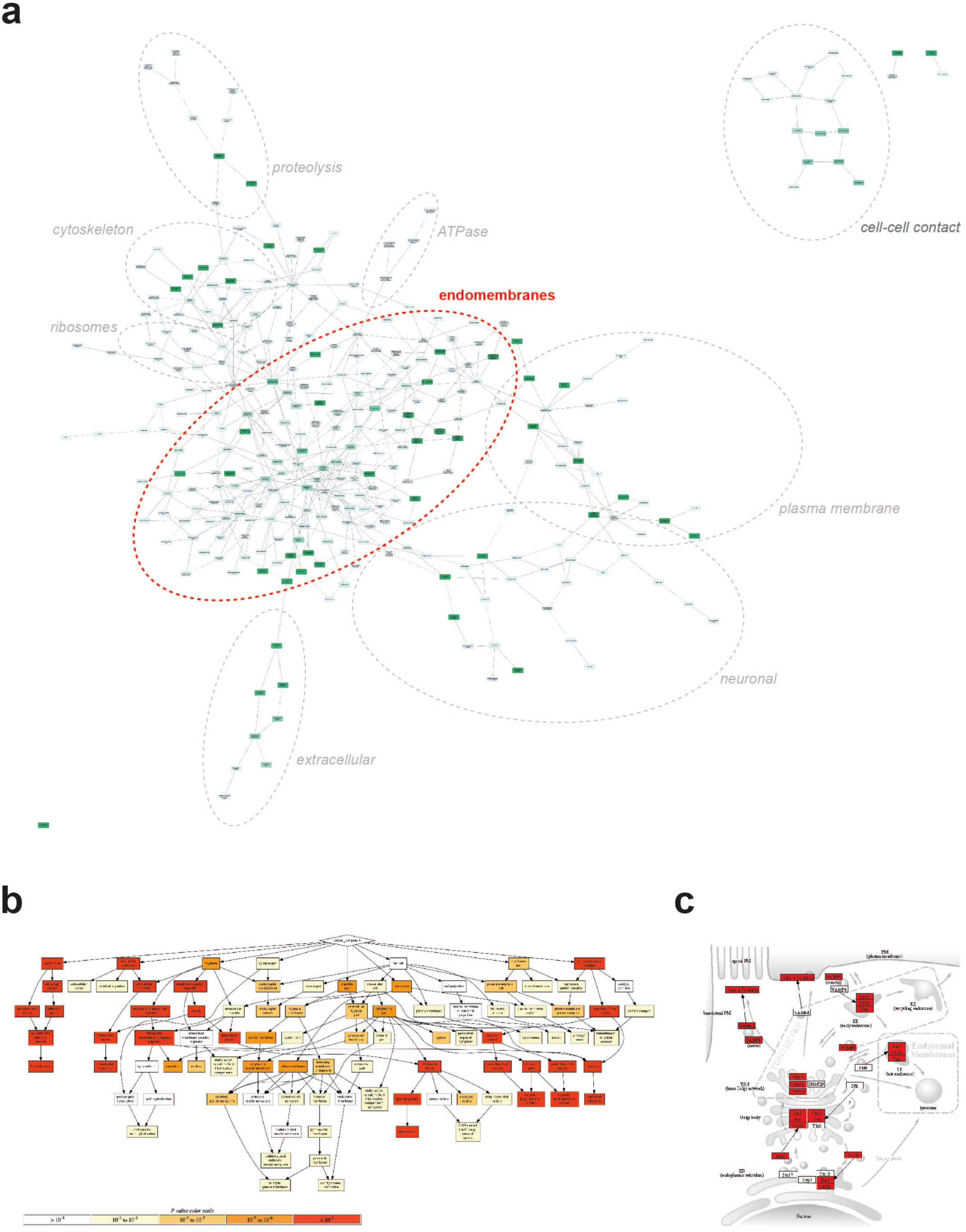
Analysis of 1710 candidate proteins identified in mass spectrometry analysis of Vangl1-positive sucrose gradient fractions. **(a)** Network representation of cellular components using GOnet (https://tools.dice-database.org). Note that the majority of proteins identified can be annotated to endo-membranes. **(b)** Flowchart representation of cellular components via Gorilla (https://cbl-gorilla.cs.technion.ac.il). Again, endo-membranes are prominently represented. **(c)** KEGG pathway analysis using ShinyGO0.80 (http://bioinformatics.sdstate.edu/go/) identifies most genes involved in endomembrane sorting.

**Supplemental Figure S7:**
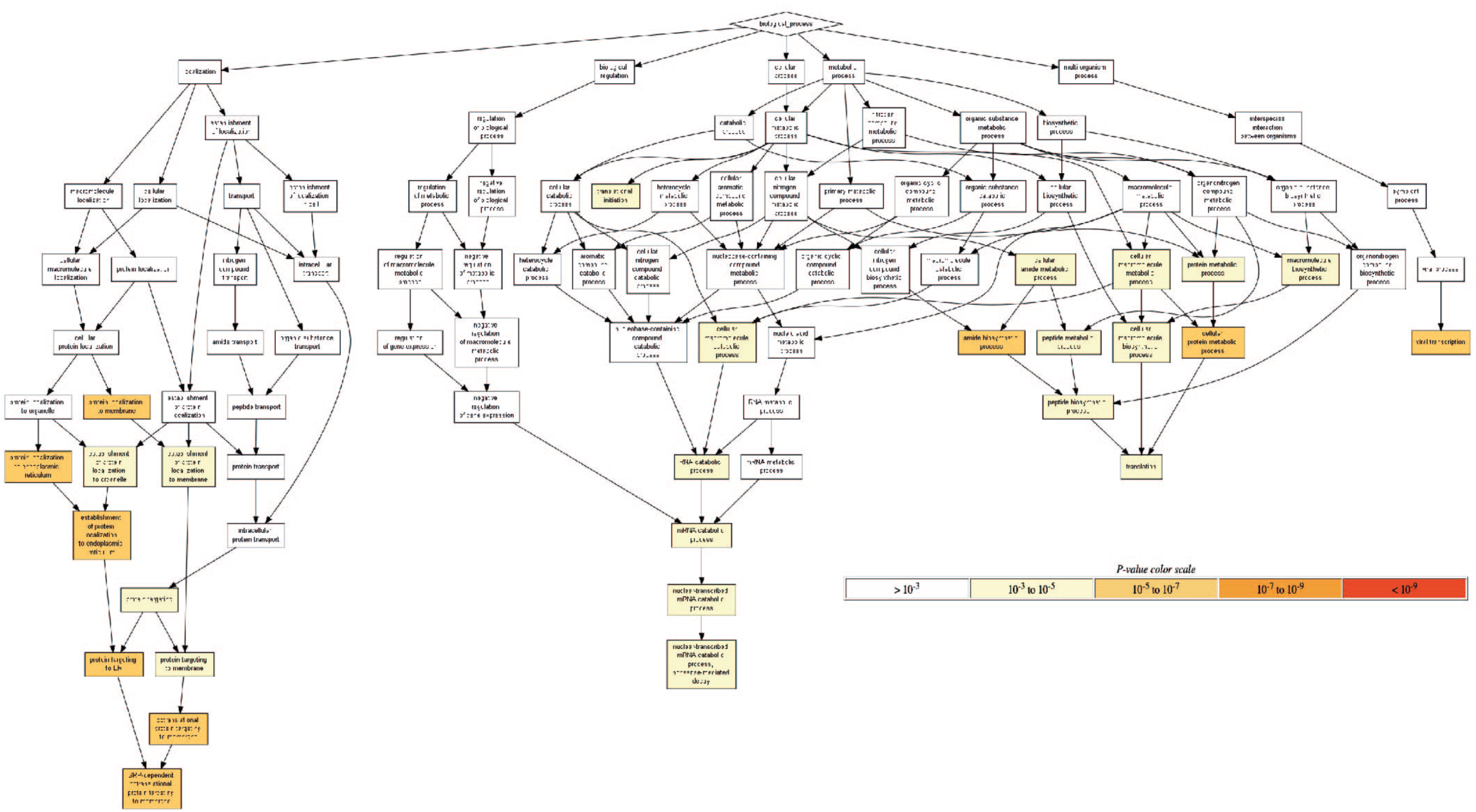
Analysis of 312 candidate proteins identified in mass spectrometry analysis of Vangl1 pulldown in the presence and absence of flow. Flowchart representation of cellular processes via Gorilla (https://cbl-gorilla.cs.technion.ac.il). (N = 5 biological repeats, 3 without and 2 with flow).

**Figure S8:**
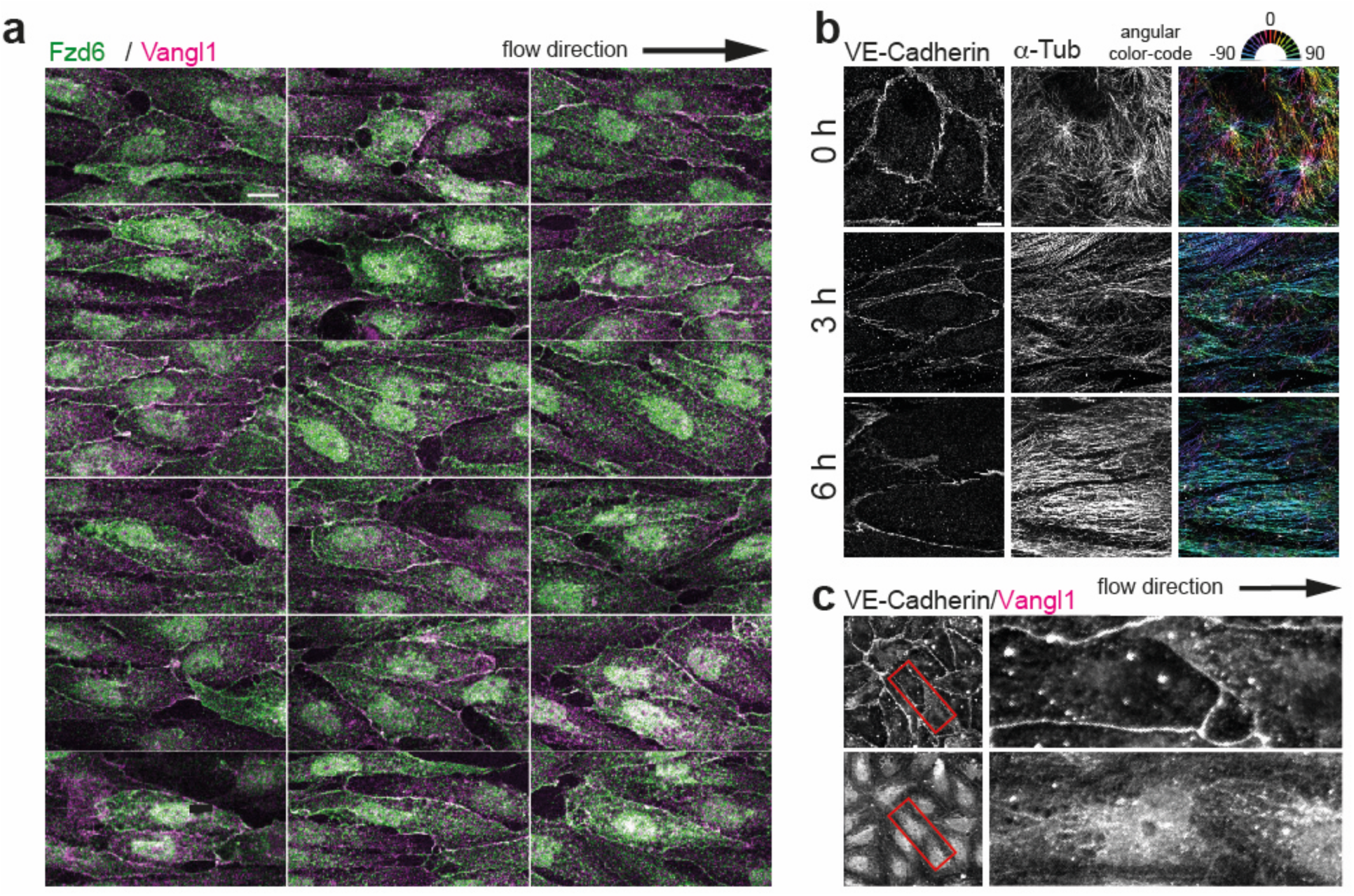
Localization of endogenous core PCP proteins and microtubules in HUVECs. **(a)** Representative images of HUVECs exposed for 24 hours the shear forces show Fzd6 (green) and Vangl1 (magenta) at opposite sides of the cell. **(b)** Upon flow induction, microtubules align along the flow axis that correlates with the longest cell axis. Microtubules are color-coded in an angle-dependent fashion. **(c)** Vangl1 appears in lines along the longest cell axis. Cells were fixed and stained with antibodies against VE-Cadherin and Vangl1. Scale bars (a,b,c), 10 μm.

**Figure S9:**
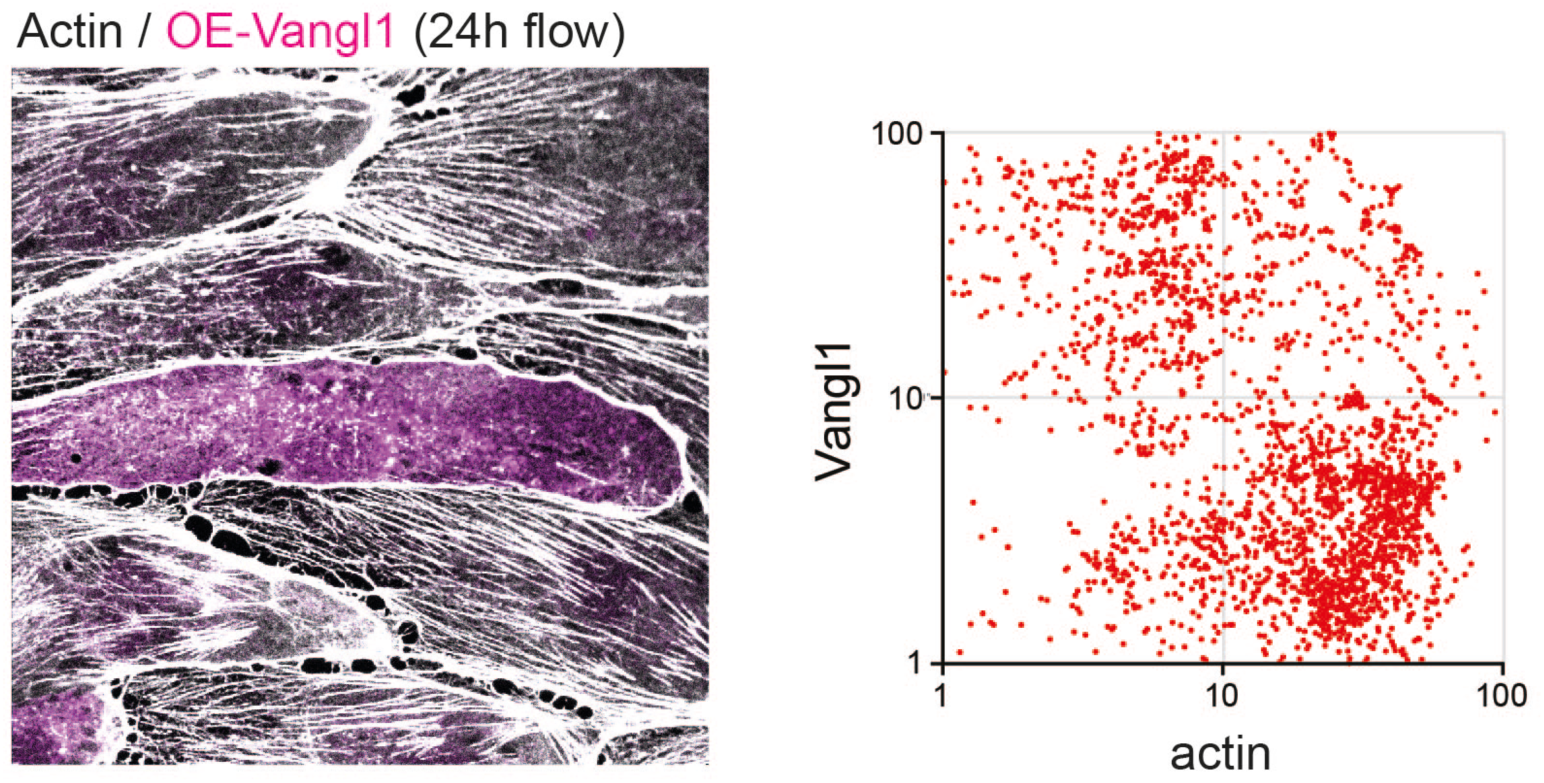
Overexpression of Vangl1 reorganizes actin filaments in HUVECs. HUVECs were transfected with plasmids expressing Vangl1, incubated for 24 hours, and then fixed and stained with fluorescently labeled markers against the FLAG tag (magenta) and actin (white). From left to right, a single cell and a scatter plot depicting Vangl1 vs. actin levels are shown (N = 5, n = 2410).

**Figure S10:**
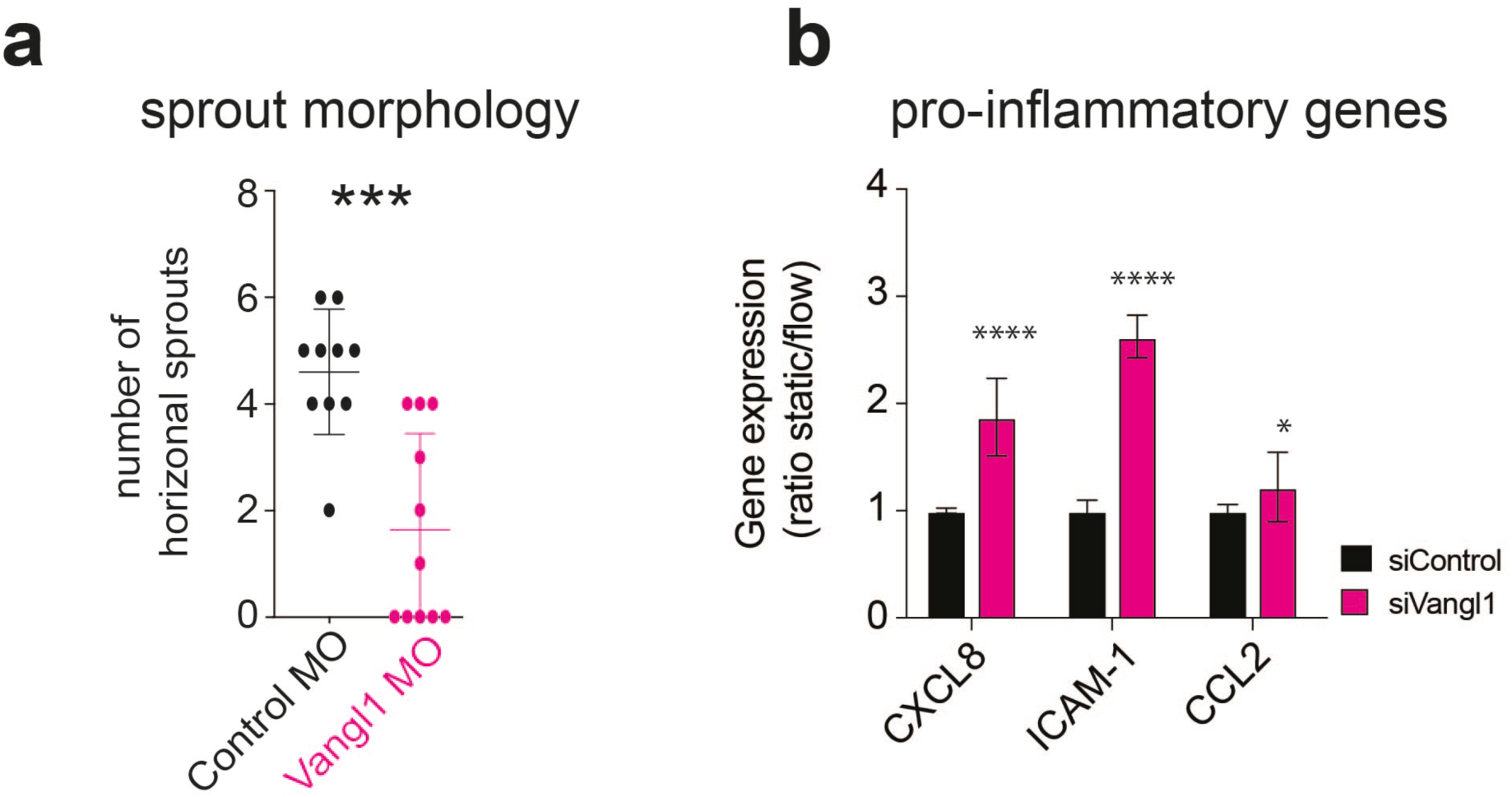
Knockdown effects of Vangl1 in developing zebrafish embryo and HUVECs. **(a)** Loss of Vangl1 leads to reduced horizontal sprouting in developing zebrafish embryos. *Vangl1* morpholinos were injected into embryos, and tail vein formation was analyzed at 48 hpf. Embryos lacking Vangl1 displayed fewer horizontal connections. (Control morpholinos: N = 3 repeats, n = 10 embryos; Vangl1 morpholinos: N = 3 repeats, n =11 embryos). **(b)** Loss of Vangl1 leads to an increase in pro-inflammatory gene expression in HUVECs. Total RNA was isolated 24 hours after transfection with siRNA directed against Control (black) or Vangl1 (magenta). Real-time PCR shows a significant increase in the expression of the pro-inflammatory genes Cxcl8, Icam-1 and Ccl2 (N = 3 biological repeats).

### SUPPLEMENTARY TABLES

**Table S1:**
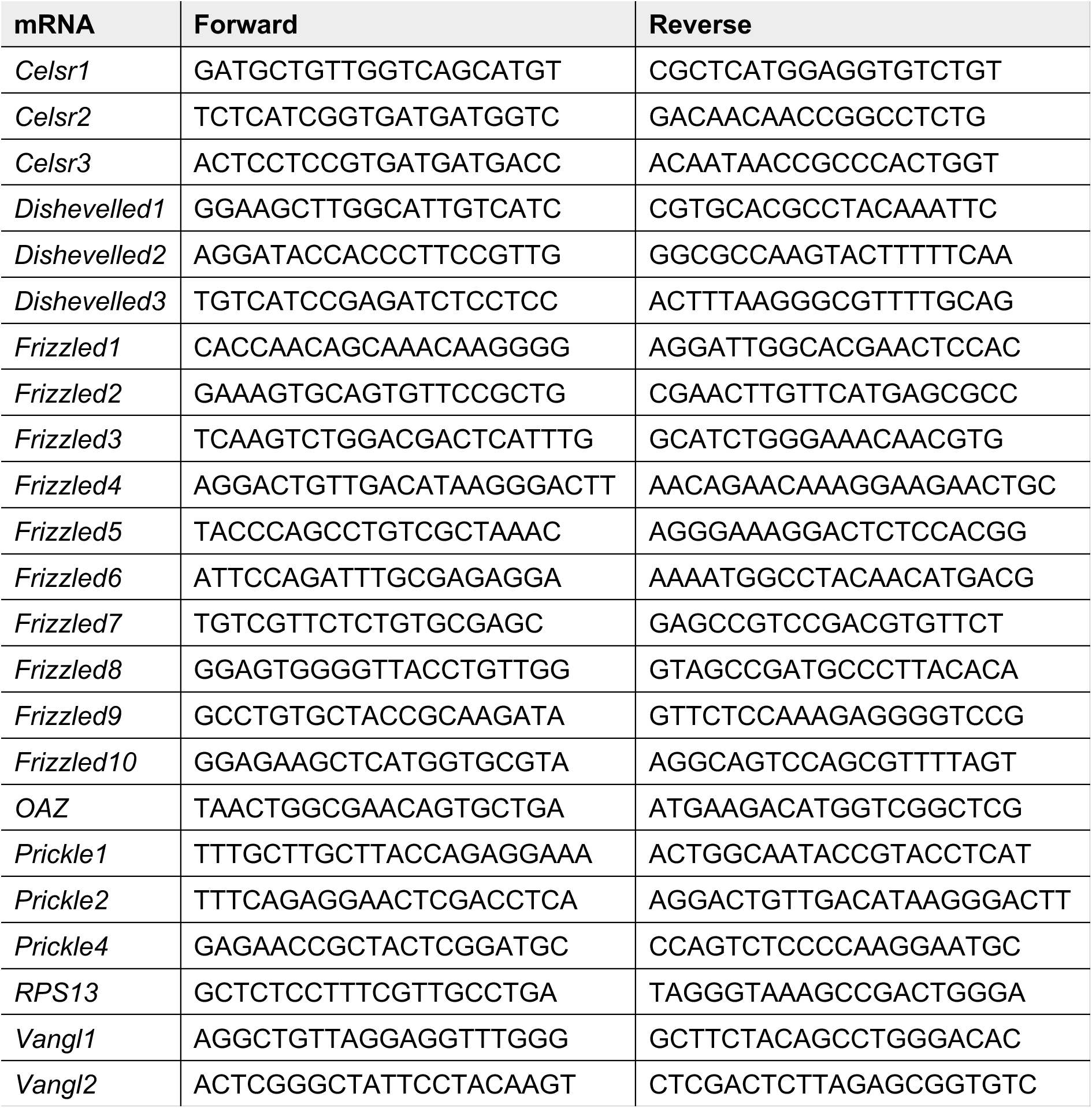
Primers used to validate PCP gene expression in HUVECs.

**Table S2:**
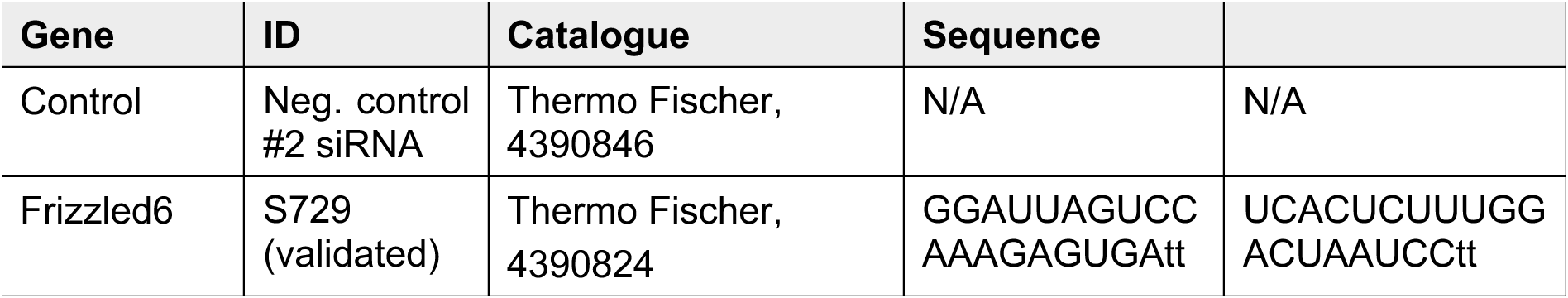
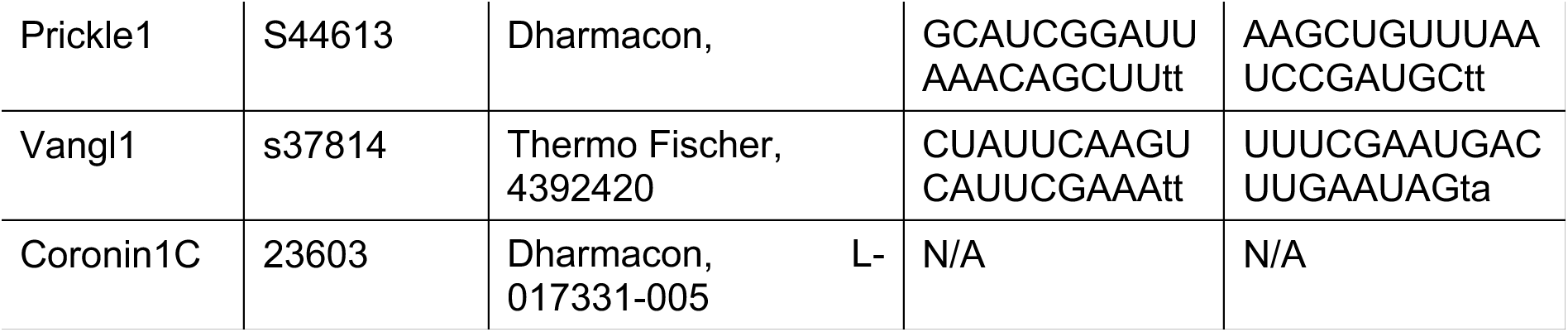
siRNA sequences used to knock down PCP mRNA levels in HUVECs.

**Table S3:** Antibodies used to probe protein levels (Western blot, WB) and localization (Immunocytochemistry; ICC) in HUVECs.

| Antibody | Cat Nr. | Concentration |
| --- | --- | --- |
| anti-Vangl1 (rabbit) | Sigma-Aldrich, HPA025235 | WB 1:1000; ICC 1:50 |
| anti-Vangl1 (mouse) | Sigma-Aldrich, Amab90600 | WB 1:1000; ICC 1:50 |
| anti-Prickle1 | Santa Cruz, sc-393034 | WB 1:500 |
| anti-VE-cadherin (rabbit) | Abcam, ab33168 | WB 1:1000; ICC 1:200 |
| anti-Coronin1C | Santa Cruz, sc-130448 | WB 1:1000; ICC 1:200 |
| anti-VE-cadherin (mouse) | Santa Cruz, sc-9989 | WB 1:1000; ICC 1:200 |
| anti-VE-cadherin (goat) | Biotechne, AF938 | WB 1:1000; ICC 1:200 |
| anti-Frizzled6 | Sigma-Aldrich, HPA017991 | WB 1:1000; ICC 1:200 |

### SUPPLEMENTARY LISTS

**List S1:**
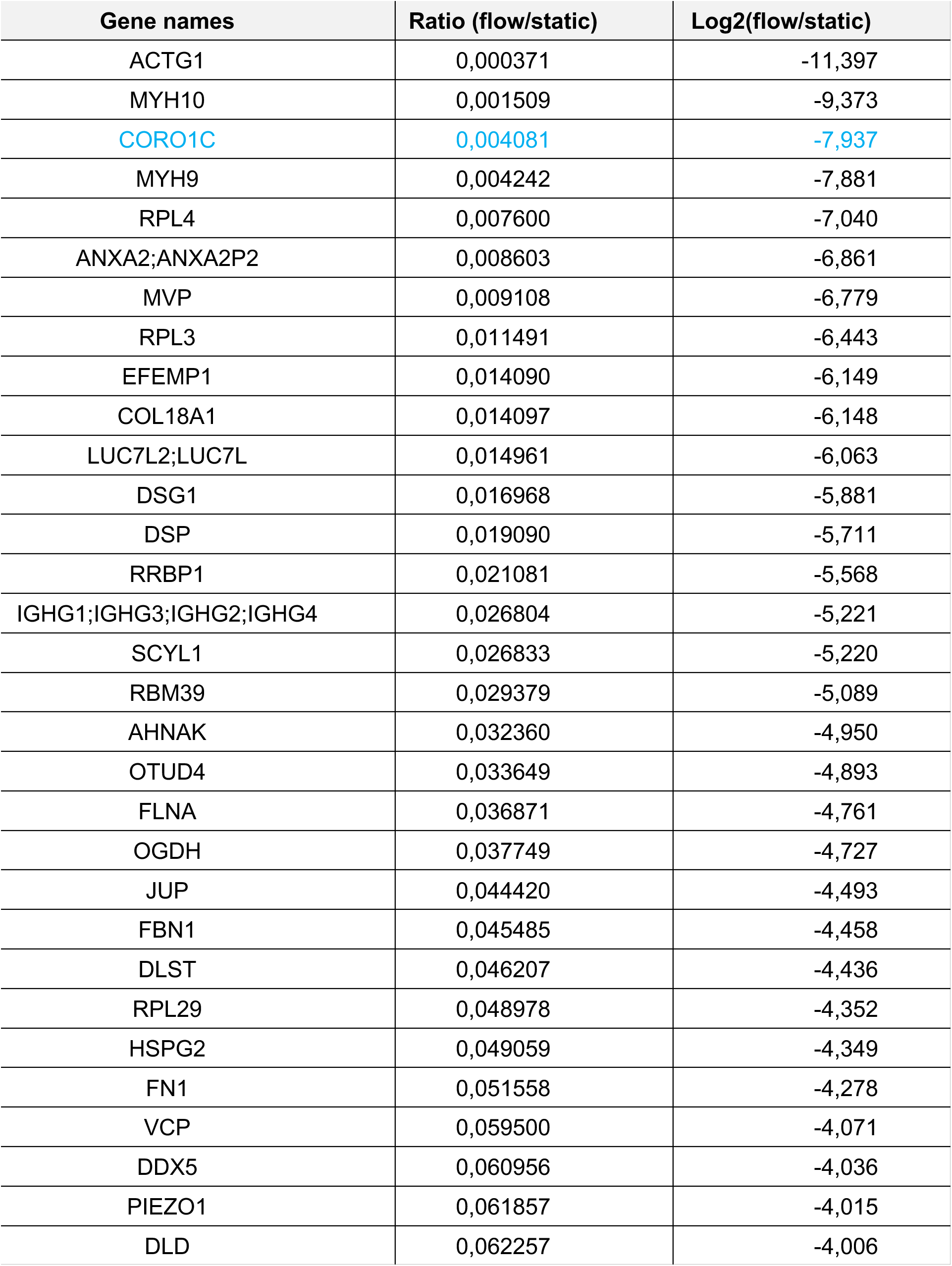
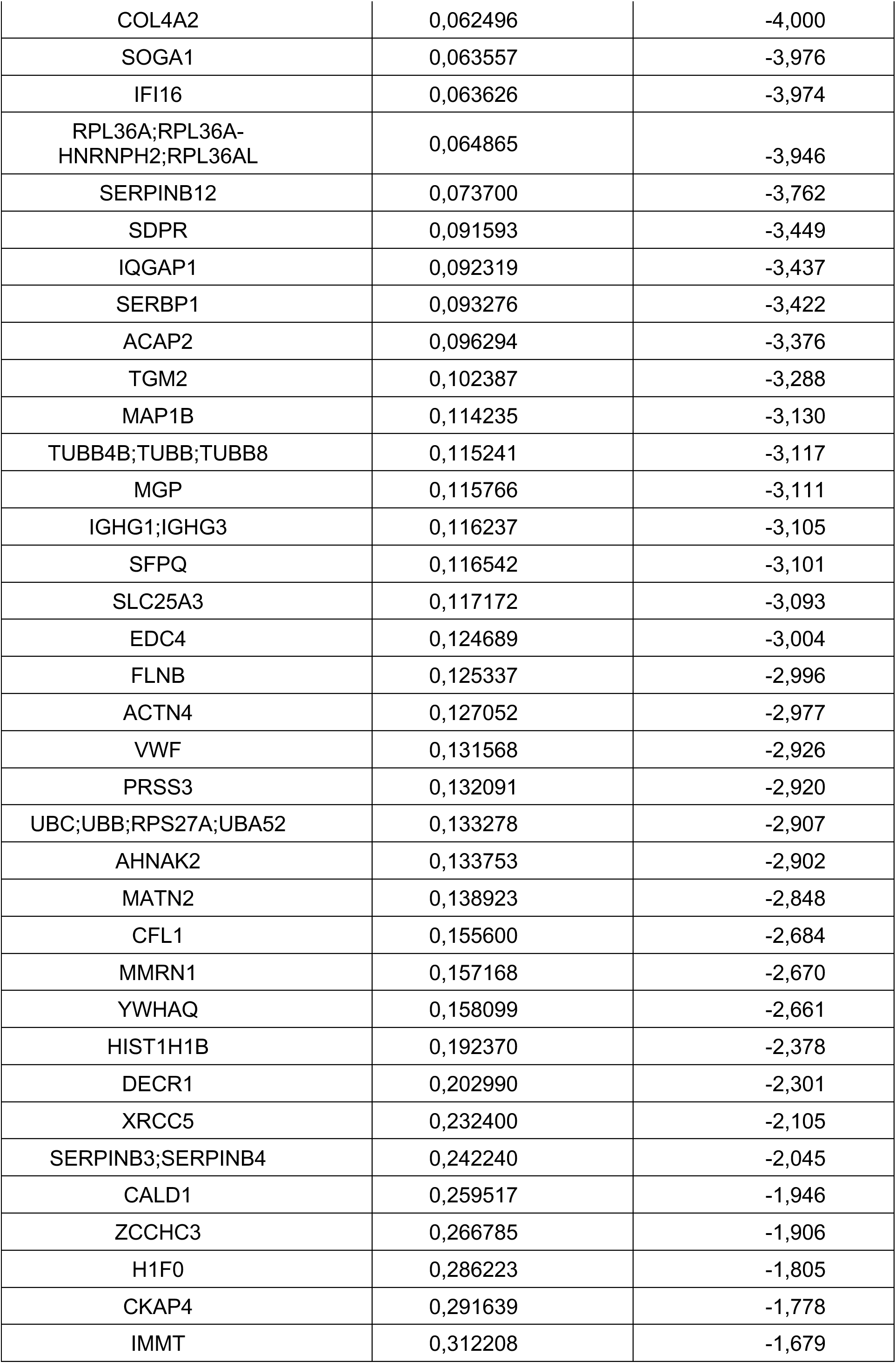
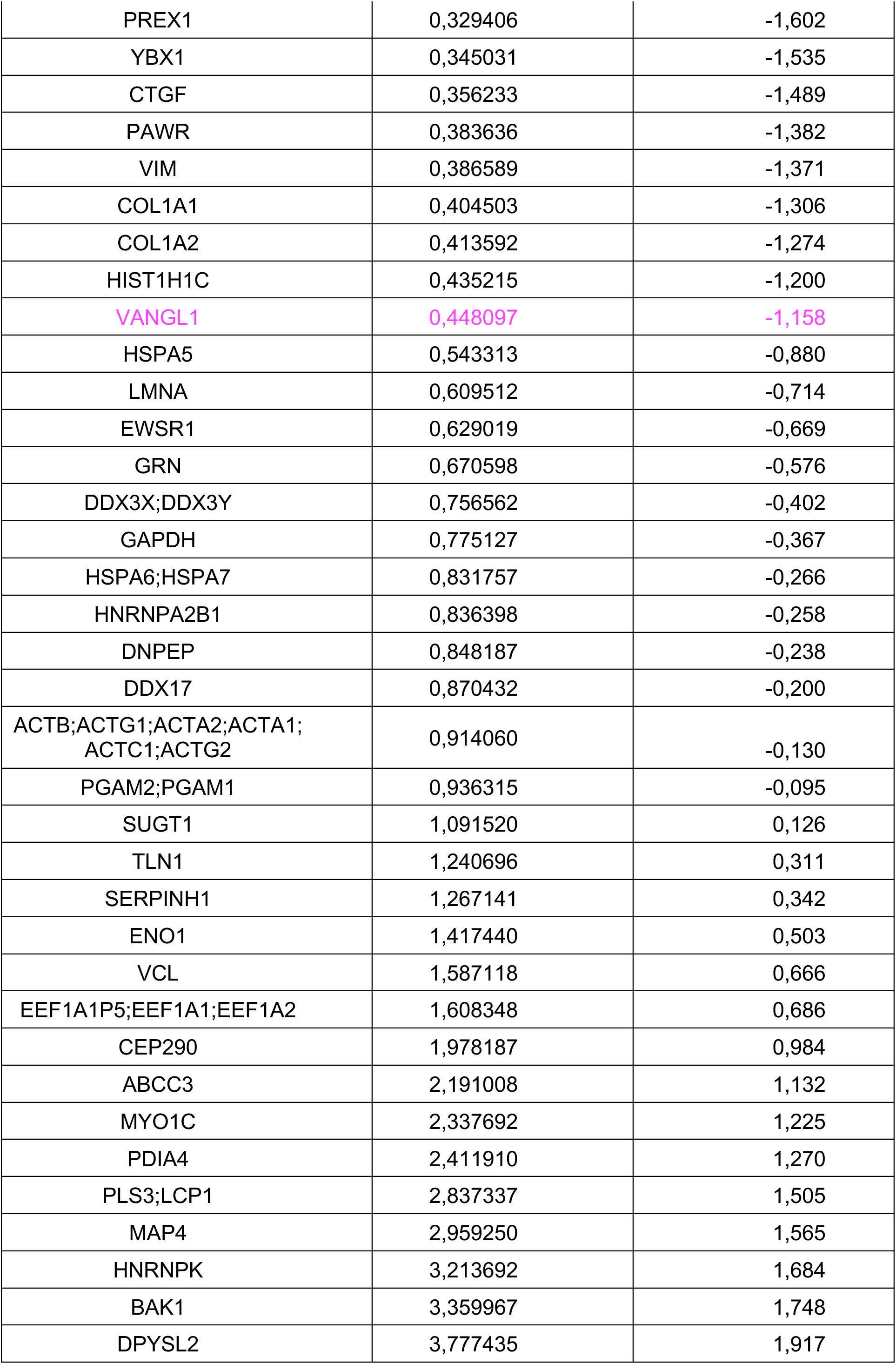
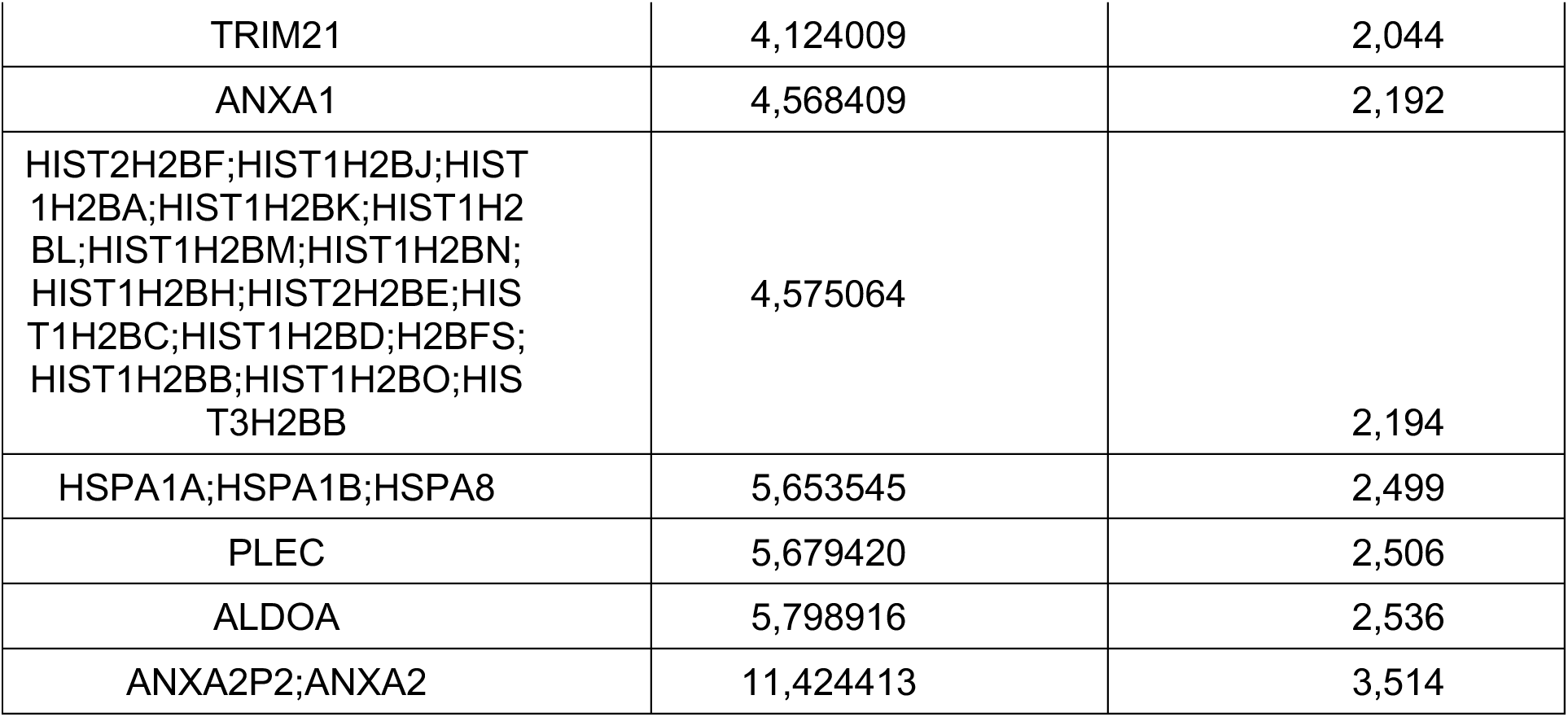
Proteins identified in MS analysis of Vangl1 pulldown to display differential expression upon exposure to shear forces.

## Notes

### Competing Interest Statement

The authors have declared no competing interest.

